# Representations of the intrinsic value of information in mouse orbitofrontal cortex

**DOI:** 10.1101/2023.10.13.562291

**Authors:** Jennifer J. Bussell, Ryan P. Badman, Christian D. Márton, Ethan S. Bromberg-Martin, L.F. Abbott, Kanaka Rajan, Richard Axel

**Affiliations:** Department of Neuroscience, Columbia University; New York, NY, 10027, USA; Zuckerman Mind Brain and Behavior Institute, Columbia University; New York, NY, 10027, USA; Department of Neurobiology, Harvard Medical School; Boston, MA, 02115, USA; Kempner Institute, Harvard University; Cambridge, MA, 02138, USA; Department of Neuroscience, Washington University School of Medicine; St. Louis, MO, 63110, USA; Howard Hughes Medical Institute; Chevy Chase, MD, 20815, USA

## Abstract

Animals are motivated to seek information that does not influence reward outcomes, suggesting that information has intrinsic value. We have developed an odor-based information seeking task that reveals that mice choose to receive information even though it does not alter the reward outcome. Moreover, mice are willing to pay for information by sacrificing water reward, suggesting that information is of intrinsic value to a mouse. We used a microendoscope to reveal neural activity in orbitofrontal cortex (OFC) while mice learned the information seeking task. We observed the emergence of distinct populations of neurons responsive to odors predictive of information and odors predictive of water reward. A latent variable model recapitulated these different representations in the low-dimensional dynamics of OFC neuronal population activity. These data suggest that mice have evolved separate pathways to represent the intrinsic value of information and the extrinsic value of water reward. Thus, the desire to acquire knowledge is observed in mice, and the value of this information is represented in the OFC. The mouse now provides a facile experimental system to study the representation of the value of information, a higher cognitive variable.

## Main Text

Humans and other vertebrates exhibit an innate drive to acquire information, even if it does not lead to extrinsic reward. Experiments to understand information seeking have largely involved paradigms in which subjects choose to receive information that reveals but does not alter future uncertain outcomes (*1–9*). Subjects are even willing to pay for this information, sacrificing reward for information of no apparent extrinsic value (*10–14*). These experiments suggest that the acquisition of information can be of intrinsic value (*15, 16*).

Rewards such as food or water activate physiological pathways that lead to neural representations of reward value (*17–19*). Information, however, is a higher-order cognitive variable whose value representations have been more elusive. In functional MRI studies in humans, information seeking tasks elicit a BOLD signal in midbrain and prefrontal reward-associated structures (*10, 20–25*). Electrophysiological recordings in information seeking paradigms in monkeys have identified a cortex-basal ganglia circuit that extends from the anterior cingulate cortex (ACC) to the lateral habenula, a major regulator of midbrain dopaminergic cells (*1, 26, 27*). Neurons in this pathway respond when reward outcomes are uncertain and diminish their response upon the acquisition of information that eliminates this uncertainty. Moreover, midbrain dopamine neurons respond to stimuli predictive of both information and liquid reward, reflective of prediction error in classical mechanisms of reinforcement (*16*). This suggests a circuit in which information activates a cortex-basal ganglia loop, eliciting a dopamine-mediated reward prediction error.

Predicted value is consistently observed in orbitofrontal cortex during decision making tasks in both primates and rodents (*28–35, 35, 36*). In information seeking tasks, single recorded neurons in monkey orbitofrontal cortex (OFC) exhibit a preference for either predicted information or external reward, suggesting the presence of distinct representations of intrinsic and extrinsic value (*11*). However, the representation of the intrinsic information value predicted by a sensory stimulus in neural populations and the process by which it acquires value through reinforcement in OFC remain unknown. The circuit that allows an organism to recognize that a sensory stimulus predicts information, a cognitive variable, remains elusive.

We have developed a paradigm to ask whether mice value information that resolves the uncertainty of a probabilistic reward outcome but does not offer any apparent extrinsic reward. In this task, mice learn that odor stimuli predict information, and the intrinsic value of information could be quantified by the willingness of mice to pay for information by sacrificing water reward. We have imaged populations of neurons during learning to reveal the emergence of a distinct representation of the intrinsic value of information in orbitofrontal cortex. These results imply that the mouse exhibits an innate desire to acquire information, a higher order cognitive function. Moreover, the value of information and the value of extrinsic reward are encoded by population structures that can be disentangled in the mouse OFC.

## Mice value information

We have developed an information seeking task in which water-deprived mice learn to poke their nose into a center port (Fig1A). There they receive one of two different odors that direct them either to the left or right side port (forced trials) where they receive a water reward on 25% of the trials. In the side ports, the mice are exposed to one of two different odors followed by a 10s delay prior to the receipt of water. In the Information port, one odor accurately predicts the receipt of water, whereas the second odor signals the absence of a water reward. In the other No Information port, the mice receive one of a different pair of odors that provide no information as to the outcome of the trial. Whereas the reward probability is equivalent at the two side ports, the Information port differs in that it provides a mouse with information that predicts the reward outcome. Once the mice have learned the structure of the task a third odor is also offered at the center port allowing a choice between the Information or No-Information port (choice trials) on 1/3 of trials. Thus, one set of odors at the center port *predicts* information, and a different set of odors at the information side port *provides* information. Mice are trained to perform this task in multiple stages lasting 5-8 weeks (fig. S1). A bias for choice of the Information port would imply that mice seek knowledge of no apparent extrinsic value since this information does not alter the structure of the task or the reward outcome.

**Fig. 1.**
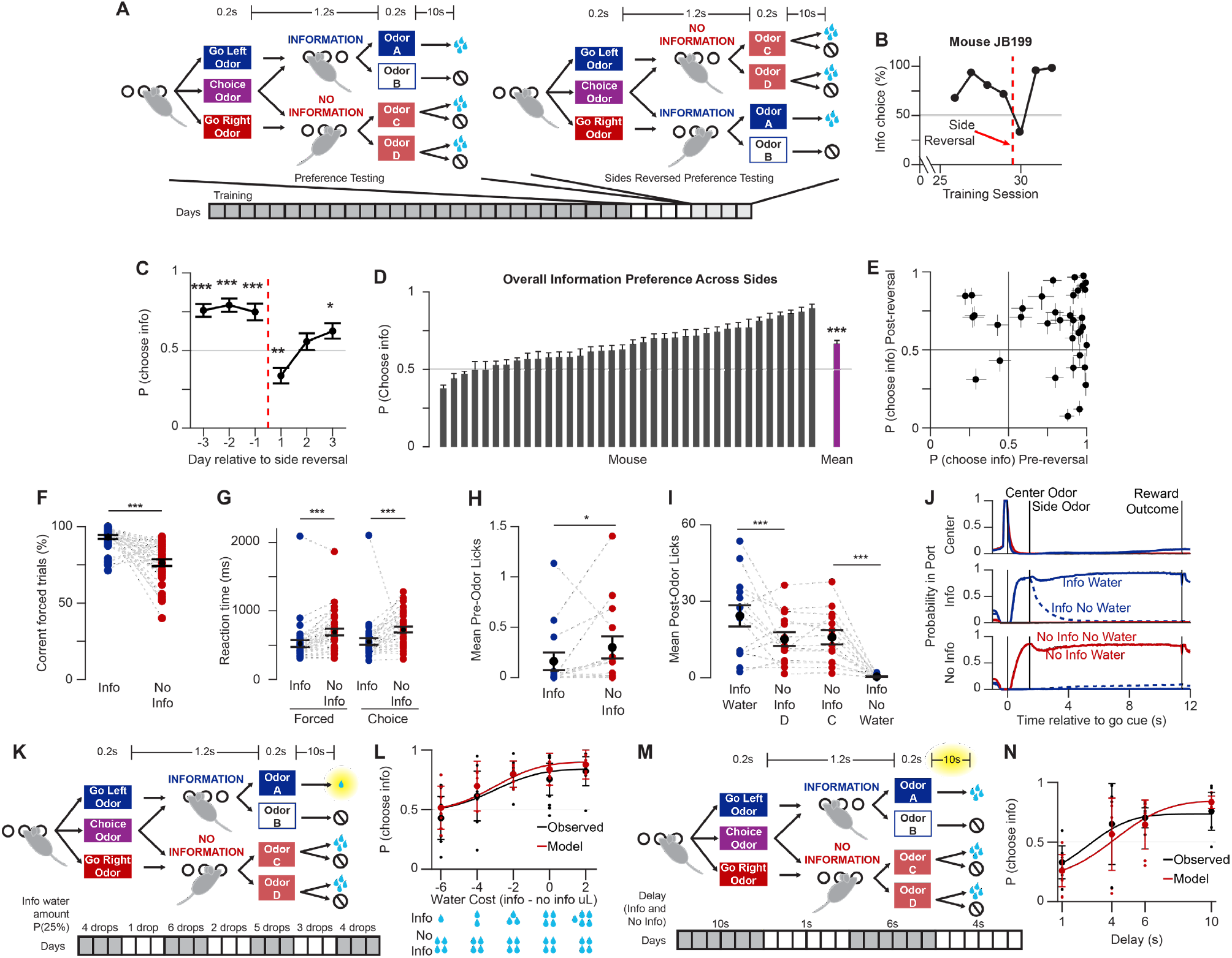
Mice value information. (**A**) Information choice task. (**B**) Information preference on free choice trials for an example mouse. Dotted line shows the reversal of the side port identities as shown in (A). (**C**) Information preference surrounding side identity reversal across animals. Red dashed line indicates reversal of left-right side identities as in A. For each session, the cross-animal mean estimated probability of choosing information (binomial fit) and 95% confidence interval is shown, Wilcoxon sign rank, N=36. (**D**) Information preference on all choice trials with information on either side. For each mouse, estimated probability of choosing information (binomial fit) and 95% confidence interval is shown. Mean shows mean and std err of estimates across animals, p<0.001 mean differs from 0.5, sign-rank test. (**E**) Preference for information before and after side identity reversal. A point for each animal shows the estimated probability of choosing information (binomial fit) and 95% confidence interval in up to the last 200 free choice trials before and after side reversal. (**F**) Error rates on forced trials. Red and blue points show mean correct choices (binomial fit) across the last two sessions prior to side reversal for each animal, black shows mean and std err across animals, Wilcoxon sign rank test. (**G**) Mean reaction times by trial type in the last two sessions prior to side reversal for each animal in blue and red. Black shows mean and std err across animals, Wilcoxon sign rank test. (**H**) Licks at the side port after entry, prior to odor delivery. Red and blue points show mean licks across the last two sessions prior to side reversal for each animal, black shows mean and std err across animals, Wilcoxon sign rank test. (**I**) Licks between side odor delivery and time of reward outcome. Red and blue points show mean licks across the last two sessions prior to side reversal for each animal, black shows mean and std err across animals, Friedman’s ANOVA with tukey-cramer correction for multiple comparisons. (**J**) Probability of mouse nose poking in each port by trial type, N=36. (**K**) Schematic of water value tradeoff sessions. (**L**) Probability of choosing information (binomial fit) at different information port water amounts, N=6. Black points, single animal preference in last two sessions at each value; bars, cross-animal mean +/-95% CI. Red, predictions from RL decision model. (**M**) Schematic of delay manipulation sessions. (**N**) Probability of choosing information at different delay durations, N=6. Plot as in L.

When given a choice at the center port between visiting the Information or No Information side ports mice chose the Information port on 78% (mean, 71-83%, 95% confidence interval, p<0.001) of the free choice trials across several sessions of preference testing (fig. S1I). We performed reversals of the Information side for each animal and observed that mice maintain their preference for the Information port despite its new location (Fig. 1, B-E, fig. S1J). Several additional observations demonstrate that mice indeed learn the structure of the task and the information-predictive meaning of the center port odor cues. On forced trials mice enter the Information port 170ms faster than they enter the No Information port, with fewer errors when choosing the Information port (Fig. 1, F and G, fig. S1L). We also recorded licks at the reward spout. Mice began licking the water spout as soon as they entered the No Information port but withheld licks in the Information port until the informative odors were presented (Fig. 1H), indicating that they learned that the Information side port odors reveal whether they will receive water on that trial. The water reward rate did not differ between Information and No Information on correct trials, and mice displayed consistent information preference across the duration of each training session rather than water satiety affecting their choices (fig. S1, K and L). These observations indicate that the mice understand the predictive meanings of the three odors at the center port and distinguish the two side ports by their information content rather than a difference in water reward.

We also observed behaviors at the side ports that reflect conditioned responses to the water reward value predicted by each odor cue (*5, 9*). Following odor A, the cue that predicts water at the Information port (100% reward), animals remained in the port and licked, but following odor B, the cue signaling the absence of water at the Information port, the mice did not lick and left the port (0% reward; Fig. 1, I and J). Following the two odors at the No Information port (25% reward), mice remained in the port and licked (Fig. 1, I and J). We considered the possibility that leaving may be of value to a mouse, and this might contribute to their preference for the Information port. However, we observed that after odors that signal the absence of water at the Information port, mice do not roam freely in the box. Rather, they remain near the ports, actively engaged in the task (fig. S2, A-D). Moreover, we developed a separate version of the information seeking task that provided a cue signaling the reward outcome 2s before it occurred on all trials. In this version of the task, we observed that mice leave both the Information and No Information side ports on all trials prior to receiving reward but return to the port in proportion to their expectation of water reward (fig. S2, E-H). Mice also display a strong information preference in this version of the task (fig. S2F, N=8, 81% mean information choice, p<0.01).

The preference we observe for the Information port presumably reflects the value of information. We therefore asked whether mice were willing to pay for information, a signature of information value (*10–13, 37*). We varied the amount of water reward on information trials over blocks of several sessions (Fig. 1K). Mice continue to prefer information even at the expense of diminished water reward, which allowed us to quantify information value as equivalent to 4-6uL of water (Fig. 1L, black).

In our information seeking task, mice move from the center port to a side port where they experience a 10s delay before receiving a water reward. We expect that if we shorten this delay, the value of information should be reduced because the shortened delay may diminish the duration of the experience of either pleasurable knowledge or aversive uncertainty (*2, 5, 20, 38*). We have therefore manipulated the value of information in this task by varying the length of delay on both sides, Information and No Information, across blocks of sessions (Fig. 1M). We observed that the preference for information decreases with decreasing delay (Fig. 1N, black).

We developed models to predict choices to receive information across sessions in which we changed the water reward value and delay (See Methods, fig. S3). We fit a Rescorla-Wagner (*39, 40*) reinforcement learning model to the choices of mice during the above water tradeoff and delay tradeoff sessions. The best-fitting model included two separate prediction error value functions, one for water and one for information, that capture the relationship between information preference, information choice history, and water reward (Eq. 9, fig. S3). We also fit a delay-related discount exponent for the value of information for each animal to quantify the degree to which delay modulates information preference (Eq. 7). This model fit the choices of all mice in these sessions to ∼70-80% accuracy (Fig. 1, L and N, red). Thus, our decision modeling analysis supports the conclusion that mice compute the value of information separately from the value of water in a manner that is dependent on the duration of the delay between information and reward.

## Representations of value in OFC during information seeking

In our behavioral task, different odors predict the intrinsic value of information and the extrinsic value of water reward. In the mouse olfactory system, the identity of an odor is represented in piriform, and this representation is largely unaltered when an odor is afforded value (*41*). Piriform projects strongly to OFC, which exhibits a neural representation of value only after odor learning (*41*). We have therefore asked whether we can detect neural representations of both intrinsic and extrinsic value in OFC as mice perform our information seeking task. If information is of intrinsic value, the center port odors that result in movement to the Information port, whether forced or choice, should serve as conditioned stimuli (CSs) *predicting* the acquisition of information (Fig. 2A, right). At the Information side port, both odors also *provide* information. Odor A predicts water reward, whereas odor B predicts its absence. Both odors are therefore unconditioned stimuli (USs) for information. In addition, the odors at the No Information side port and odor A at the Information side port predict water reward with differing probabilities and are therefore CSs for water. We have asked whether we can detect each of these representations of intrinsic information and extrinsic water values in OFC in response to the odors in our task.

**Fig. 2.**
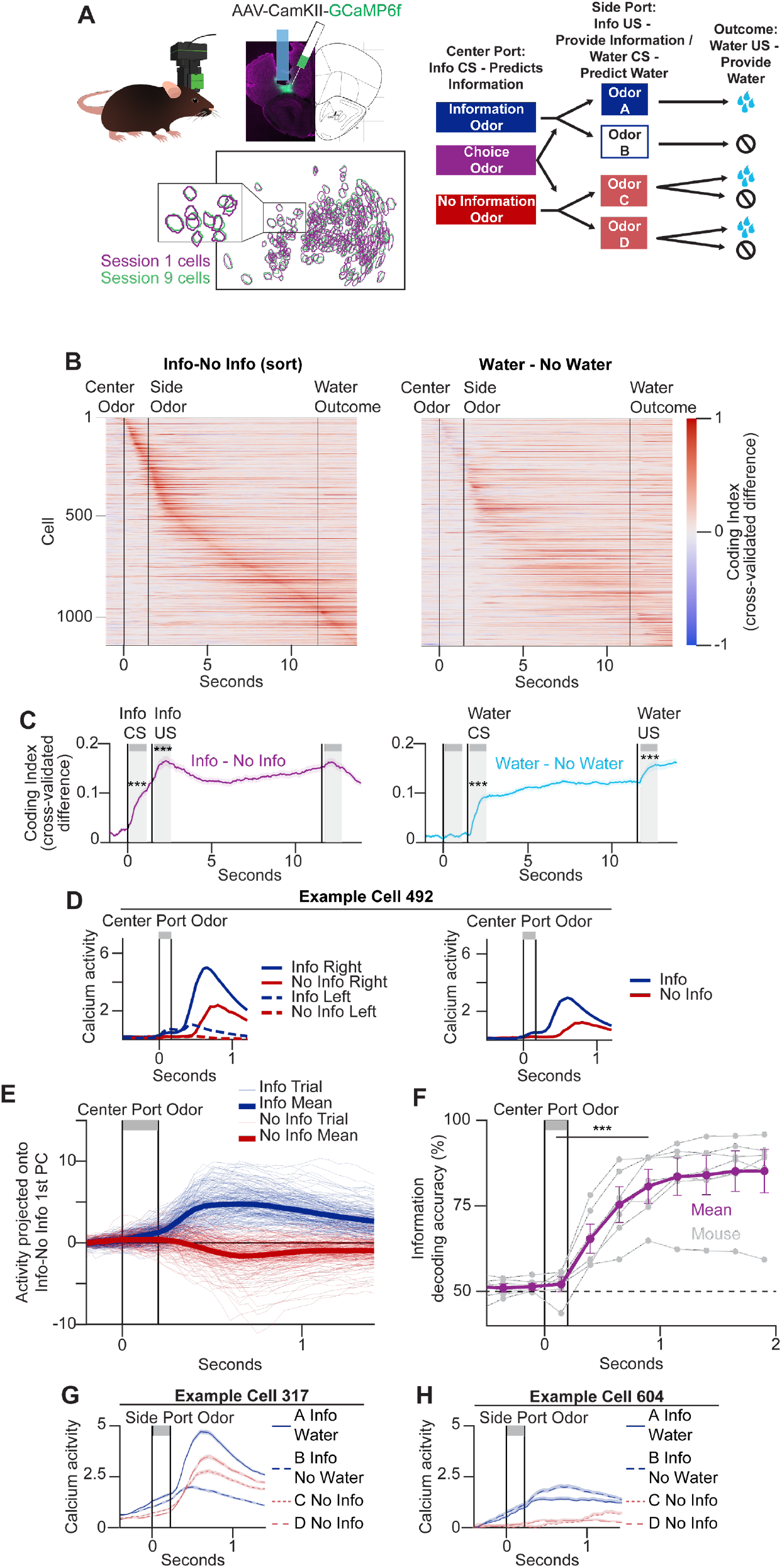
Representations of value in OFC during information seeking. (**A**) Imaging schematic. Left, miniature microscope imaging and cell registration. Blue outline shows GRIN lens implantation location in a coronal slice from an example brain. Bottom left, example field of view with cells registered across two sessions 8 training days apart. Right, schematic of odor representations. (**B)** Left, information value coding index across the population. Right, water value coding index. Cells sorted in both by information coding index (**C**) Left, population mean information value coding index. Right, population mean water value coding index. Shading shows standard error. p<0.001 difference between mean index in 1s after event and 1s before event is greater than data shuffled 1000 times. Gray shading shows 1s analyzed post-event to determine CS and US value representations. (**D**) Calcium activity of an example cell at center port odor presentation of an example cell. Left, mean calcium activity by information and side location Right, mean calcium activity across all information versus no information trials. (**E**) Projection of trial activity onto the first principal component of the absolute difference between mean information and mean no information center port odor responses across cells, N=1138 cells. (**F**) Accuracy of decoding of held out forced information versus forced no information trials with a linear classifier. Purple, mean +/-95% CI across mice, gray=individual animals. Significance tests whether mean accuracy greater after center port odor than before, Wilcoxon sign rank. (**G**) Mean calcium activity at the side port odor by trial type for one example cell. Shading shows std err. (**H**) Mean calcium activity at the side port odor for a second example cell.

We recorded calcium signals reflecting neural activity in 1138 neurons in OFC using miniaturized microscopes in seven mice. We registered cell identities across four sessions for each mouseincluding sessions in which we reversed the Information side port location (Fig. 2A, fig. S4, A and B). We expect that the representations of information and water value in our task may be complex. Neurons in mouse OFC represent task variables including stimulus identity, motor action, confidence, task context, and reward value (*29–33, 42–47*). Moreover, OFC neurons display mixed selectivity; individual cells encode multiple variables, and the representations of variables are overlapping^55^. We therefore computed coding indices for information and water value across the components of the trial (Fig. 2, B and C, fig. S4C). The information coding index is the mean difference in activity between information and no information trials, with the trials split into two arbitrary halves and the sign of the difference in each half taken from the other half to counterbalance the direction of the difference (Fig.2, B and C left). We similarly computed the coding index for water using the difference between rewarded and unrewarded trials (water coding index, Fig. 2, B and C right).

We examined these coding indices across the recorded OFC populations throughout trials in our task (Fig. 2, B and C, fig. S4C). We observed increases in the information coding index following the center and side port odors and in the water coding index following exposure to the side port odor and the receipt of water reward (Fig. 2C). We observed an increase in the information coding index across the population following the center port odor, reflecting the information CS (p<0.001, fig. S4E), and an increase in the information coding index upon exposure to the side port odors reflects the information US (p<0.001, fig. S5F). We also observed an increase in the water coding index following the side port odor across the population, reflecting the water CS (p<0.001, fig. S4F), and following the receipt of water, reflecting the water US (p<0.001, fig. S4G). There was little overlap between the cells comprising the CS versus US representations, for both information and water value, in accord with observations that OFC encodes CSs and USs distinctly (fig.S4, H-M, p>0.05, chi-square) (*11, 41*). These OFC responses therefore provide neural representations of a CS and US for the intrinsic value of information and a CS and US for the extrinsic value of water reward.

We observed an increased information coding index (information CS) at the center port in 19% of OFC neurons (fig. S4, D and E). Individual cells displayed mixed selectivity and responded to both predicted information and side location (Fig. 2D, fig. S5, A,E, and F) (*43, 48*). We therefore demixed the encoding of information prediction within the center port odor response by projecting each trial’s activity onto the first principal component, which captures 80% of the variance, of the difference between information and no information activity at this task epoch (Fig. 2E). The separation of activity on information versus no information trials projected along this single mode implies that the differential coding index we have computed accurately reflects encoding of the prediction of information at the center port on each trial. Moreover, we observed similar responses to the center port odors on forced and choice information trials (fig. S5, C, D, and G). We were able to decode information versus no information trials at the center port using a linear classifier with 70% accuracy on forced trials and 68% accuracy on choice trials (Fig. 2F, fig. S5B). The responses we observe to odors predictive of information at the center port reflect the choice to receive information and not the animal’s movement (fig. S5, H and I).

The increase in both the information and water coding indices in response to the odors at the side ports reflects two different value representations (Fig. 2C, fig. S4, D and F). At the Information side port, odor A predicts water reward and is therefore a water CS+. Odor B predicts the absence of water reward and is a water CS-. However, both odor A and odor B also may elicit neuronal activity because they provide information and are an information US. Indeed, the responses to odors A and B were highly correlated across cells (r=0.73, p<0.0001, Pearson’s correlation, Fig. S5, L and K). We therefore calculated the water coding index from the differential activity between odors A and B, which reflects the water CS. Both odor A and odor B provide information, whereas the No Information side port odors C and D do not. The differential activity (information – no information trials) between them defines the information coding index at the side port and may reflect an information US.

We observed individual cells that comprise of each of these different value representations (fig. S5J). 21% of OFC cells displayed an increased water coding index at the side port (fig. S4D), reflecting the water CS, and one example is shown in Figure 2G. 25% of OFC cells displayed an increased information coding index at the side port (fig. S4D), reflecting the information US, and one example cell is shown in Figure 2H. Thus the OFC encodes both information and water value in multiple representations.

## OFC representation of predicted information scales with value

Our behavioral experiments show that as we diminish the delay between the side odors predictive of reward and the receipt of water, mice diminish their preference for information. This suggests that the value of information is reduced with diminished delay (*2, 4, 27, 49*). We therefore examined the OFC responses to odors at the center port that predict information in sessions with either a 1s or 10s delay. Across the population the number of cells exhibiting information coding was similar across both delays (45% overlap, p<0.001, chi square, fig. S6D). However, the magnitude of the information coding index for the center port odors was diminished in sessions with a 1s delay (Fig. 3, A-D). This reflected decreased responses to odors predictive of both information and no information (fig. S6, A-C). It did not reflect generally diminished OFC activity in a task with shorter delay or the passage of time (fig. S6, E-I). These data are in accord with our behavioral data and decision model that show a decreased preference for information with shorter delay and suggest that the information CS representation reflects the value of information as scaled by delay.

**Fig. 3.**
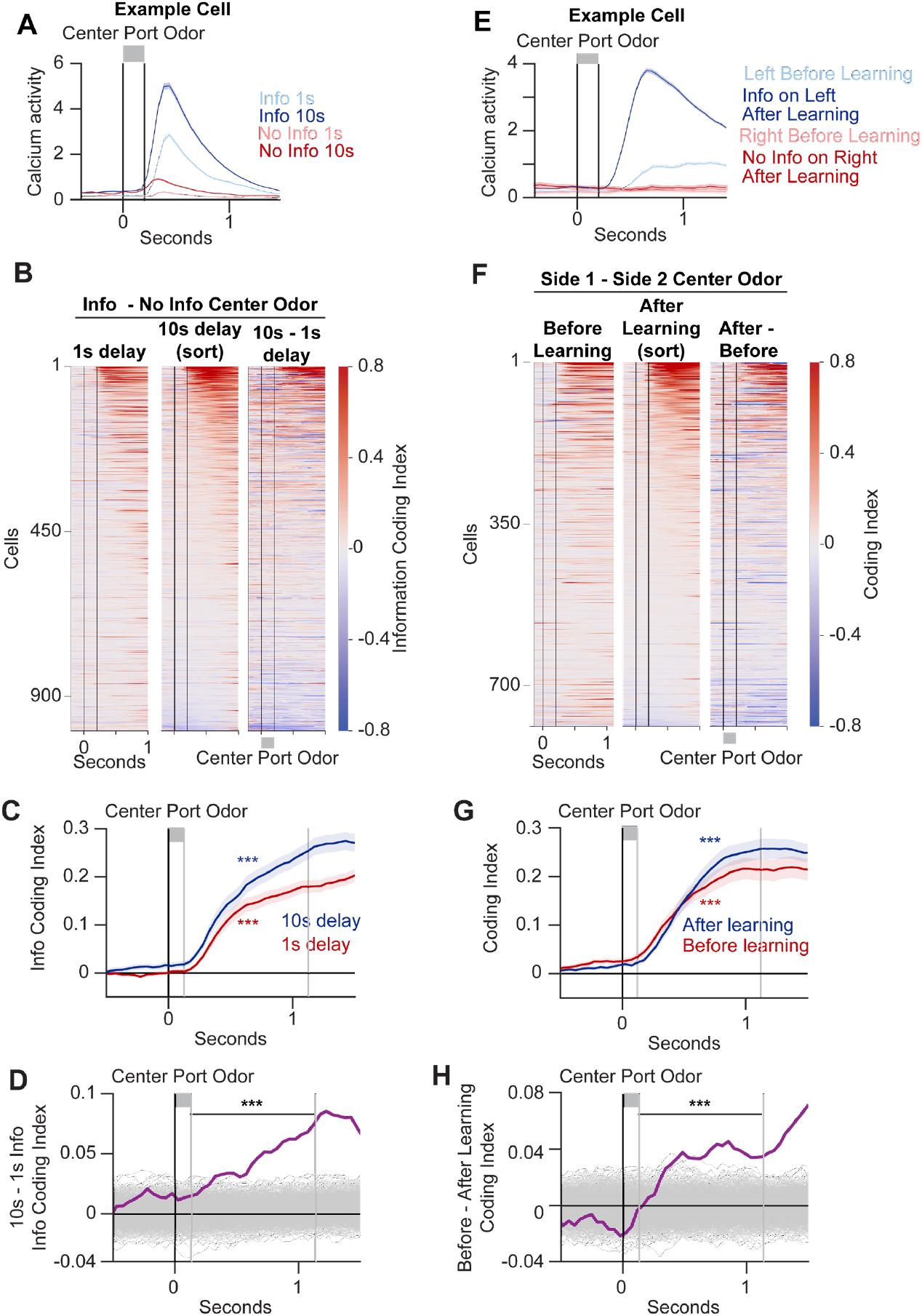
OFC representations of predicted information reflect the intrinsic value of information and emerge with learning. (**A**) Mean calcium activity by trial type for an example cell at center port odor delivery in sessions with 1s and 10s delay. Shading shows std err. (**B)** Mean coding index on forced information-forced no information trials across the population in sessions with 1s delay, 10s delay, and their difference. (**C**) Mean population information value coding index in 10s and 1s delay sessions. Shading shows std err, significance based on index difference between 1s after odor presentation and 1s before greater than shuffled data. (**D**) Mean difference between information coding index in 10s – 1s delay sessions (purple). Gray shows shuffled data. (**E**) Mean calcium activity by trial type for example cell at center port odor delivery in sessions before and after the introduction of information and probabilistic reward. Shading shows std err. (**F)** Mean coding index for each cell on forced left-forced right trials before learning and forced information – forced no information trials after learning and their difference. (**G**) Mean population coding index for left – right side in sessions before learning and information – no information in sessions after learning, plot as in C. (**H**) Mean difference between side and information coding index across sessions before and after learning (purple). Gray shows shuffled data.

## The emergence of a representation of information value with learning

We recorded the emergence of representations of the prediction of information at the center port as the mouse learned that information was available at one of the two side ports. During early training, prior to exposure to informative odor, the two different center port odors directed movement to the left or right port, where mice received equal water reward on every trial (fig. S6J). Information providing-odors were then introduced, and water was delivered with 25% probability at both ports (fig. S6J). Thus, mice learned that one port provided information as to whether they would receive water reward.

We observed increased differential activity in response to odors at the center port upon learning. We computed the coding index from differential activity at the center port between left and right side trials prior to learning. After learning, the odor that previously directed the mouse to the left port, for example, now also predicted information at the left port, whereas the odor directing the mouse to the right port now predicted no information (the information side port was balanced across mice). We computed the information coding index from the differential activity at the center port after learning. We observed that the information coding after learning is greater than the coding index between left and right prior to learning (Fig. 3, E-H). This increase is due to an enhanced response to odors predictive of information in cells that overlap with those that previously responded to the odor prior to learning (55% overlap, p<0.001, chi-square, Fig. 3E, fig. S6, K-M). Thus, a representation predictive of information emerges from the population of cells that previously represented odor directing the mouse to a particular side to receive water reward.

## Distinct representations of intrinsic and extrinsic value

The differential neural activity in response to odor predictive of information at the center port reflects a CS for the receipt of information, whereas the differential activity in response to odor predictive of water at the Information side port reflects a CS representation for water reward. We asked whether the CS representation of information is distinct from the CS representation of water. We compared the population of cells exhibiting significant coding of odor predictive of information at the center port (CS info) with the population of neurons with significant coding of odors that predict water (CS water) at the side port (Fig4A). These two CS representations, comprising 19% and 21% of OFC cells respectively, are distinct and exhibit only 23% overlap with each other (Fig. 4B, p>0.05, chi-square). Moreover, there is no correlation between the coding indices for the two CSs across the OFC population (Fig. 4C, r=0.04, p=0.1, Pearson’s correlation). This suggests that information and water value prediction are encoded by largely separate ensembles of OFC cells.

**Fig. 4.**
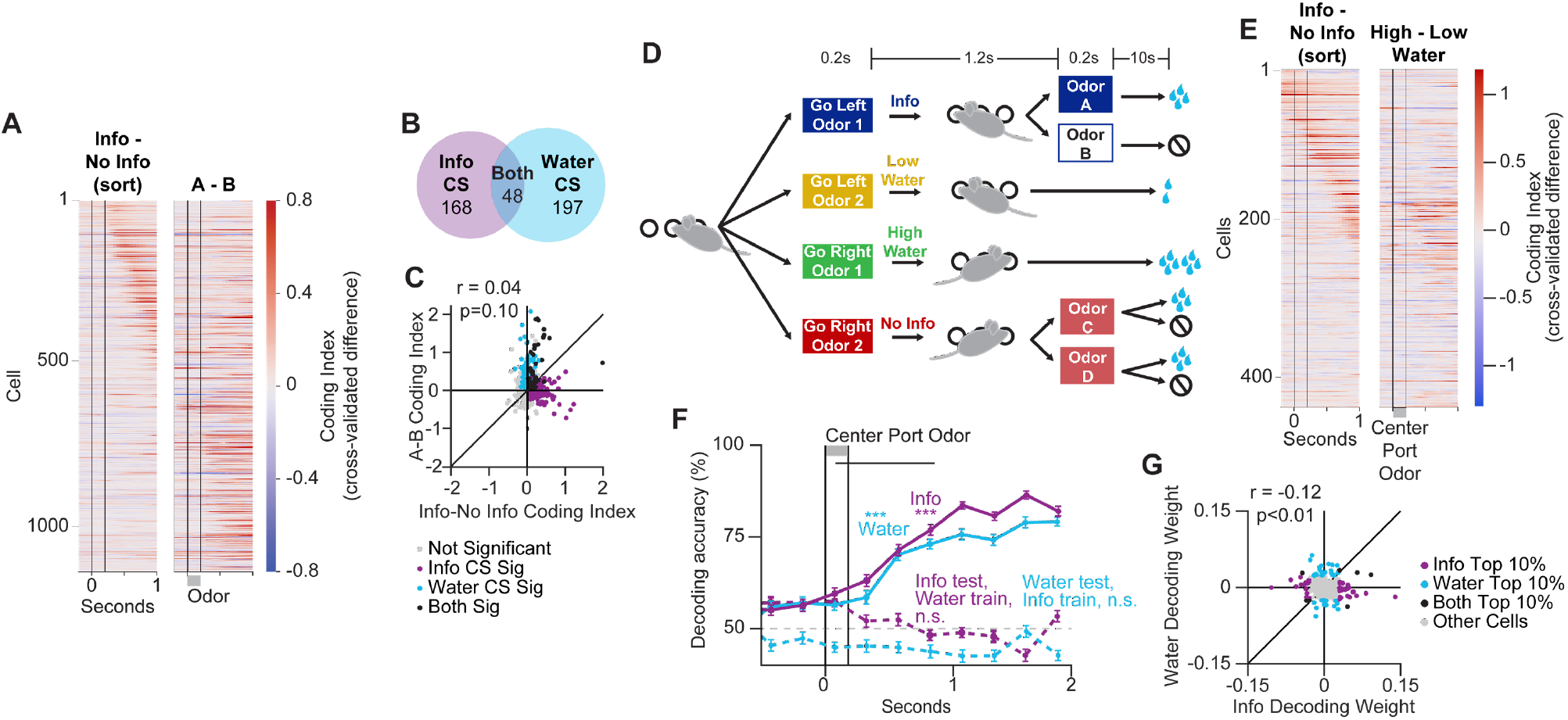
Representations of intrinsic and extrinsic value in OFC are distinct. (**A**) Information CS and water CS coding indices for each cell. Left, information value coding index (forced information – forced no information trials) at the center port odor. Right, water value coding index (odor A (water) -odor B (no water)) at the Information side port odor. (**B**) Number of cells with differential activity (significant coding index increase after odor) between the information and no information forced trial center port odors and between the information side port odors A and B. N=1138, overlap, p>0.05, chi-square. (**C**) Pearson’s correlation between the mean information coding index in 1s following the information and no information forced trial center port odors (information CS) and the water coding index in response to the information side port odor A (water) and B (no water) (water CS). (**D**) Task providing four center port odors that predict information, no information, large water reward, and small water reward. (**E**) Mean information (left) and water (right) coding indices at the center port odor across the population in the 4 center odor task. (**F**) Decoding held out information vs. no information trials (purple) and water large vs. small amount trials (cyan) at the center port odor with classifiers trained on either matched data (solid lines) or data from the other trial types (dashed lines). Significance tests whether accuracy significantly increases across 1s after the center odor (horizontal black line) more than shuffled data. (**G**) Correlation between information and water decoding weights. Weight assigned to each cell in the linear classifier for information vs. no information (x-axis) or large vs. small water (y-axis) trials. Cells colored by the absolute value of the decoding weight in the top 10% for each decoder. Pearson correlation shown.

The information CS and water CS representations are evoked at different epochs within our task, and we considered that the distinct representations we observed of the predicted value of information and water might reflect this temporal separation (*50*). We therefore designed a task in which the representations of the predicted value of information and water could be analyzed at the same task epoch. We trained mice that had previously learned the information seeking task on a modified task in which four odors were presented at the center port. As in the original task one odor directed the mouse to the Information side port and a second to the No Information port. Two additional odors were presented, one signaling a large water amount at one side port and a second a small water amount at the other side port, each delivered with 100% probability (Fig. 4D). Mice displayed faster reaction times for information versus no information and large versus small value water reward trials, indicating they understood the meaning of the center port odors in this task variant (fig. S7A). We imaged OFC activity before and after reversal of the side locations in this task (fig. S7E). We computed the information coding index and a coding index for large versus small water amount (Fig. 4E, fig. S7, B and D). We observed that distinct subsets of cells encoded the information CS and the water CS in response to odors at the center port (fig. S7C, p>0.05, chi-square). We then trained two different linear classifiers, one to decode information versus no information trials and one to decode large versus small water amount trials at the center port (Fig. 4F, mean accuracy 69%, p<0.001, for information, 65% for water, p<0.001). The classifier trained to decode information did not accurately decode water value and vice versa (Fig. 4F, mean accuracy 42% for decoding water with information classifier, 50% for decoding information with water classifier). Moreover, these classifiers used weights that were weakly negatively correlated across cells (Fig. 4G, r=-0.12, p<0.01, Pearson’s correlation), indicating that the cells contributing most to the decoding of information value were different from those contributing to the decoding of water value. We asked whether we could identify subgroups of cells that determined the decoding of information versus water. We selected the top 10% of cells in each classifier, information and water, by the absolute value of their weight within each classifier (Fig. 4G). We then trained new classifiers using either only these cells, only other cells, in the absence of these cells, or in the absence of other cells. We observed significantly greater accuracy in classification of information vs. no information trials when the top information coding cells were included, and greater accuracy in classification of large vs. small water amount trials when the top water coding cells were included (fig. S7, F and G). These data support the existence of distinct representations of odors predictive of information and odors predictive of water reward in the same task epoch.

## Representations of intrinsic and extrinsic value in low-dimensional population dynamics

We identified representations of intrinsic information and extrinsic water value at particular epochs in the information seeking task. We also asked whether we could identify these value representations as latent variables in the dynamic unfolding of activity in OFC throughout the task. We fit a latent variable model called CEBRA (Consistent EmBeddings of high-dimensional Recordings using Auxiliary variables) (*51*) to derive task-relevant low-dimensional embedding spaces by jointly fitting neural data and behavioral labels across animals (Methods, Fig.5, A-I, fig. S8). We observed that the differences between information and no information and reward and no reward trials could be decoded from low-dimensional CEBRA manifold embeddings (Fig. 5, J-L, fig. S8). We also modeled OFC activity across sessions with 1s versus 10s delays between information and outcome and observed decreased distance between the CEBRA embeddings of information and no information trials in sessions with 1s delay (fig. S8, M-O). The ability to decode information versus no information and reward versus no reward trials from the CEBRA model fit concurrently across all animals suggests that these two kinds of value are represented in OFC neural dynamics on a low-dimensional manifold during performance of the information seeking task. These analyses further support the idea that mice compute and maintain separate representations of information and water value when seeking information about water reward.

**Fig. 5.**
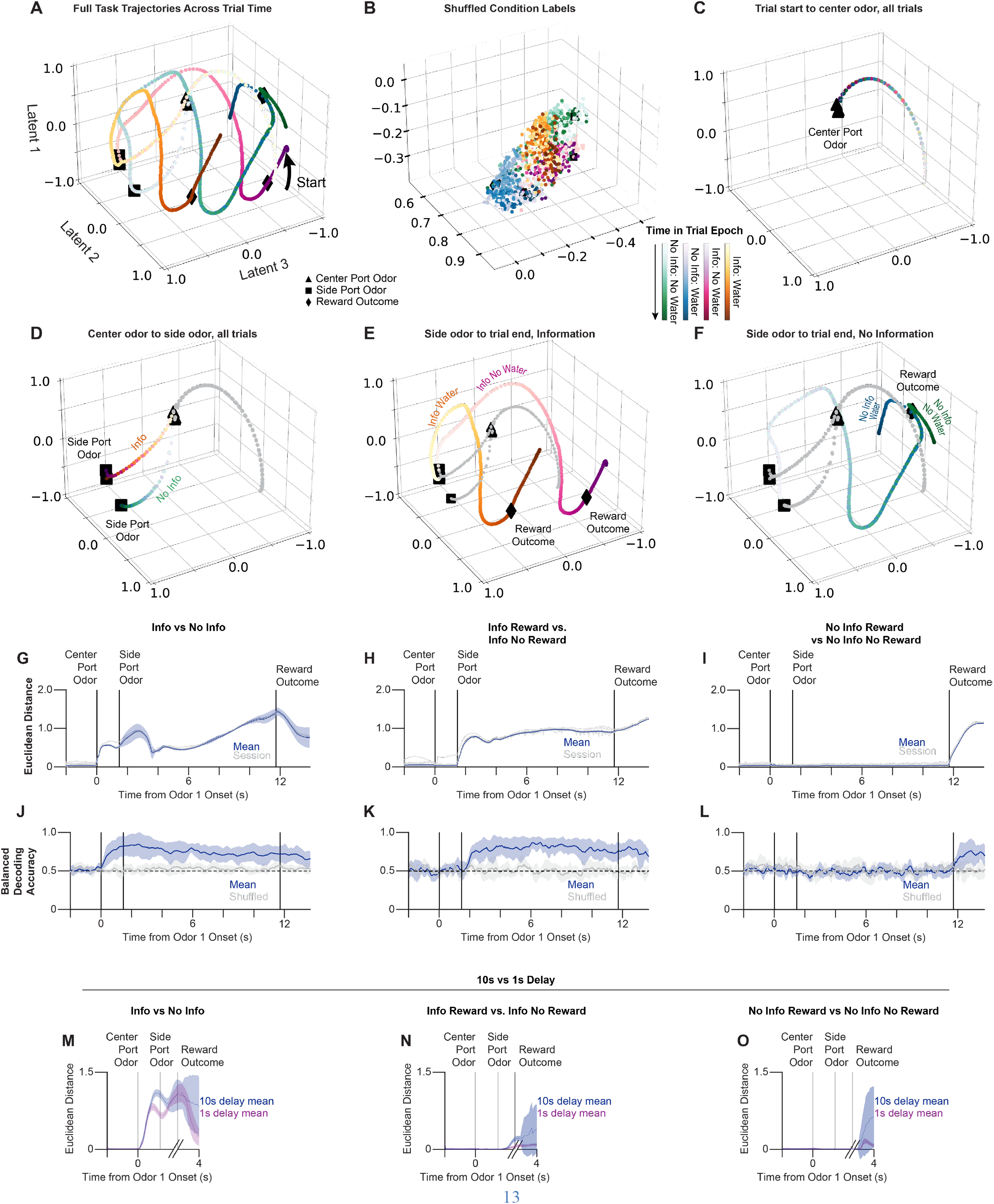
CEBRA model reveals dynamic manifold encoding of information and water value. (**A**) Plot of the first three latents of CEBRA embedding across trial time (from light to dark) for interacting labels of information and reward in a single session for a single animal. (**B**) The same plot for data with maximally shuffled labels (see Methods for details). (**C-F**) CEBRA embeddings by trial epochs (points shaded from light to dark to indicate time in trial epoch) as indicated. (**G-I**) Quantitative analysis of the embedding latents’ coordinates was then performed using Euclidean distance calculations between each type of behavior-labeled trajectory at each frame over trial-time: (**G**) the distance between information and no information trajectories, and between rewarded and unrewarded trajectories for (**H**) information trials and (**I**) no information trials. (**J-L**) To test the quality of these embeddings, as well as to find at which trial periods representations of behavioral variables were encoded in the OFC, decoding accuracy was then calculated and compared between decoding trial variables (information versus no information, and reward-information interactions), from CEBRA manifold embeddings. (**N-O**) Euclidean distance as in G-I were also calculated in a joint model that fit both 1 s delay trials and a truncated version of 10 s delay trials with ∼9 sec of the delay removed to align the first 1 s of the delay and reward periods between each delay category. Error bars in CEBRA decoding and distance plots show the 95% confidence interval.

## Discussion

We have developed a behavioral paradigm that demonstrates that mice choose to acquire information even though this information does not alter the reward outcome of the task. Moreover, mice are willing to sacrifice water reward for information, suggesting that in mice, as in several other species, information is of intrinsic value (*2–6, 10, 16, 21, 22, 52, 53*). Our paradigm has enabled the study of information seeking behavior in mice, allowing the use of methods that are not available in other information seeking species such as monkeys or humans.

Longitudinal calcium imaging during the information seeking task allowed us to follow the emergence of a representation of predicted information value during learning at the level of population structure. We observed that the representation of predicted information elicited by odor reflects information value scaled by delay, which our decision model suggests mice calculate separately from water reward. Moreover, we observed distinct representations of odors predicting information value and odors predicting water value along separate decoding dimensions in both linear classifiers at specific task epochs and on a manifold within OFC dynamics. These data suggest that mice have evolved dedicated pathways in OFC that represent the intrinsic value of information and the extrinsic value of water reward. Electrophysiological recordings in monkeys during an information seeking task conceptually analogous to our paradigm similarly reveal orthogonal coding of information and water reward in single neurons in OFC (*11*). The presence of distinct representations within OFC may allow the organism to separately learn and update different elements of a rewarding stimulus and execute different behaviors in response to distinct value representations.

Representations of the value of information must involve a complex process of cognitive abstraction because recognition that an odor provides information requires knowledge of its predictive nature despite the lack of its association with an external reward. Indeed, we observed a representation of odors that provide information irrespective of the water value associated with this information. This poses the question as to how the brain initially recognizes that an odor cue provides information of intrinsic value. Models of classical conditioning involving rewards of extrinsic value, such as food or water, invoke known physiologic responses that activate subcortical brain structures that lead to the release of dopamine and downstream reinforcement (*17–19*). Analogous models for rewards of intrinsic value suggest that in our information seeking task, there must be an odor-responsive neural ensemble that encodes the value of information and is capable of reinforcing information-predictive odor cues. However, the nature of the brain structures capable of recognizing the innate value of information and activating both the cortex-basal ganglia loop and distinct OFC subpopulations remain elusive (*11, 26*). Although dopamine may reinforce multiple neural populations, one population can only be reinforced in the context of information and the second only in the context of water.

Why does an organism value information of no apparent extrinsic value? One class of models argues that information seeking is a consequence of secondary reinforcement (*5, 54–60*). The receipt of odor A (the odor predictive of water reward) at the information side port may be overvalued and secondarily reinforce the choice of information at the center port. A second model posits that odor A at the side port boosts “anticipatory delight” enhancing the value of the information port (*2*). These models may not easily account for observations that humans and other animals exhibit a desire for information about upcoming aversive events and events that have already occurred (*10, 22, 61–63*). Others have argued that the resolution of the aversion of uncertainty provides a basis for the value of information (*27, 60, 62, 64–68*). Perhaps the simplest explanation argues that information led, at some time during the evolution of a species, to extrinsic value. Indeed, in natural environments, knowledge improves an organism’s model of the world, and its acquisition improves decision making and augments survival (*38, 69*). This quest for information may therefore have been fixed by selection and now generalizes to circumstances in which the acquisition of knowledge may be of no extrinsic value. This model suggests that the desire for knowledge evolved by assigning innate pleasure to the acquisition of information. Thus, the value of information may ultimately reflect the pleasure it provides.

## Acknowledgments

The authors would like to thank Evan Schaffer for his suggestions on cell registration and motion correction. They would like to thank members of the Axel lab, Ilya Monosov, Mackenzie Mathis, and Jackie Gottlieb for helpful discussions. They would also like to thank the research assistants and students Alexander Knue, Theodore Hannah, Ashwin Viswanathan, Deniz Bingul, Adamu Awak, and Maya Sharma Campbell who performed mouse behavioral training as well as Nataliya Zabello and the staff of the Zuckerman Institute for Comparative Medicine for their expert mouse care. We would like to acknowledge Tanya Tabachnik and the Zuckerman Institute Advanced Instrumentation Core for their help with design of behavioral chambers. We would like to acknowledge the team from Stellate Communications for their help with mouse illustration. We would also like to acknowledge Adriana Nemes, Clayton Eccard, and Miriam Gutierrez for their laboratory support and assistance in drafting this manuscript.

## Funding

NIH RF1 DA056403 (R.P.B., C.D.M., K.R.)

NSF CAREER Award (K.R.)

James S. McDonnell Foundation (K.R.)

McKnight Foundation (K.R.)

CIFAR (R.P.B., C.D.M., K.R.)

Simons Foundation (J.J.B., L.F.A., R.P.B., C.D.M., K.R.)

Gatsby Charitable Foundation (L.F.A.)

Howard Hughes Medical Institute (R.A.)

## Author contributions

Conceptualization: RA, JJB, ESBM, KR

Methodology: JJB, RPB, CDM, ESBM, KR, LA, RA

Formal Analysis: JJB, RPB, CDM, LA

Investigation: JJB

Visualization: JJB

Software: JJB, RPB, CDM

Funding acquisition: RA, LA, KR

Supervision: RA, KR

Writing – original draft: JJB, RA

Writing – review & editing: JJB, RA, LA, ESBM, RPB

## Data and materials availability

Data and code will be deposited in open databases prior to the paper’s publication.

## Supplementary Materials

Materials and Methods

Figs. S1 to S8

Tables S1 to S11

References (70–81)

## Materials and Methods

### Subjects

All experimental and surgical protocols were performed in accordance with the guide of Care and Use of Laboratory Animals (NIH) and were approved by the Institutional Animal Care and Use Committee at Columbia University. Mice were wild type C57/BL6 (Jackson Laboratories strain 000664), aged 8-45 weeks, of both male and female sex. Mice were water restricted at least 2 days prior to the initiation of training and maintained at >80% of their initial body weight throughout experiments. Mice were handled by experimenters for approximately 5-15 minutes for at least two days prior to beginning behavioral training. All animals were maintained under reverse cycled 12hr light/12hr dark conditions and single housed after beginning water restriction. Experiments were performed during the dark period of the circadian cycle.

### Behavior Training and Testing

#### Information seeking task

In general, mice were trained on the information seeking task in one session for 45-100min each weekday. Training sessions lasted until mice received at least 700uL of water or stopped engaging in trials. On average, mice received 800-1400uL of water per session and received water later that day in their home cage if daily requirements were not met during training.

#### Main task

Mice were trained to perform a 2-alternative-forced-choice task in which they chose whether to receive information revealing the trial’s reward outcome (water or no water). Behavioral training and testing were performed in custom-made acrylic boxes outfitted with three nosepoke ports (Sanworks) with odor and water delivery spouts and a solenoid valve (The Lee Company, LHDB1233418H) to deliver water reward along one wall in custom sound and light attenuating enclosures. Boxes also contained a speaker to deliver audio cues (tones). Tones were controlled by a Bpod HiFi module and played through a speaker (Peerless by Tymphany XT25SC90-04) and amplifier board (AMP2X15, PUI Audio Inc.). Behavioral hardware was controlled using Bpod microcontrollers

In the information seeking task, mice initiated trials by poking an illuminated center nosepoke port where one of three trial type odor cues was provided for 200ms. Following a 50ms 4500Hz go cue tone, the center port light went out and mice then chose either information or no information by poking into either the left or right reward port to trigger its infrared sensor. Each animal was initially assigned either left or right as the information port, pseudorandomized across animals. Mice had to indicate their choice prior to the delivery of a 200ms odor cue in the chosen side port 1.2s after the completion of the go cue or the trial had to be repeated after allowing the remainder of its duration to elapse. On some training sessions, a grace period of up to 10s was provided to extend the time available for the mouse to indicate its choice. In this case, the side odor was presented as soon as the mouse entered the correct side port. If mice did not stay in the center nosepoke for the duration of the center odor delivery and go cue, they were unable to proceed with the trial and had to re-poke the center port for the full odor duration to proceed. Trials were of three types presented in pseudorandom order in blocks of 12: Forced Information, Forced No Information, and Choice. If mice chose the incorrect port on forced trials, they did not receive side port odor or reward and experienced an uncued timeout for the remainder of the full trial time and had to repeat the trial. These procedures ensured mice were unable to avoid non-preferred trial types and equalized reward rates across trial types during training.

At the side ports, on each trial mice received one of four odors for 200ms: the information port provided odor A on 25% of trials, which was always followed by water reward, or odor B on 75% of trials, which was never followed by water reward. The no information port provided odor C on 75% of trials and odor D on 25% of trials, but the water reward was determined independently and provided on 25% of no information trials. The unequal frequency of odors C and D was designed to control for possible frequency effects on the neural representations of the side port odors A and B unassociated with their prediction of water value. The delivery of the side odor was followed by a delay period of 10s in the main task. A 4000Hz 200ms tone then indicated the outcome time on all trials, although water was only provided at that time on rewarded trials. Water rewards in the main task were 16uL provided as 4-4uL drops 50ms apart. Unrewarded trials provided a time delay to match the duration of water delivery. The reward outcome period was followed by a 3s inter-trial interval. Mice were required to be present in the chosen port for water to be delivered, but they were not otherwise required to be in the port after indicating their choice.

All odors used in the task were monomolecular neutral odorants diluted in mineral oil, through which compressed medical-grade air was bubbled in custom-built olfactometers using mass flow controllers (Aalborg, GFCS-010201) and solenoid valves (The Lee Company, LHQA1221220H), whose opening was controlled by the Bpod system valve modules. Each odor bottle had a valve located before and after it to ensure accurate odor delivery timing. An odor line of 333mL/min was combined with a carrier line of 333mL/min to deliver a total air flow rate of approximately 667mL/min into each of the three nosepoke ports. Air flow was maintained at the same rate but passed through a mineral oil control when odor stimuli were not being delivered. Odor identities were randomized across animals among four odors able to be delivered to the center port and four odors able to be delivered to either side port. Latch valves (The Lee Company, LHLA1221211H) switched odor delivery across the left and right sides. Odors were obtained at >98% purity from Sigma-Aldrich. Odors and concentrations used were as follows, at the center port: isoamyl acetate (1:10), pinene (1:5), benzaldehyde (1:10), limonene (1:5); at the side port: ethyl butyrate (1:20), acetophenone (1:5), octanal (1:5), cis-3-hexen-1-ol (1:10).

Behavior and olfactometer hardware were controlled by Bpod microcontroller systems (Sanworks, Finite State Machine r2.5) and their Matlab interface as well as custom Matlab code (Mathworks). Behavioral timestamps, video recording, and imaging frame acquisition were synchronized using external data acquisition systems (National Instruments, USB-6002).

#### Behavioral Training

Mice were trained to perform the information seeking task over several weeks of behavioral shaping progressing through the following stages.

1. **Covered Side Training (fig. S1A)**. Mice performed alternating blocks of trials where they had to trigger the center port and then enter the left or right port where they received water reward. The incorrect port was covered to prevent entry for the duration of each block of ∼50 trials (200uL total reward) each. Trials provided 1-2 drops of 4uL of water, with the reward on the two sides being equal. The center port odors that would direct mice to each side on later forced trials were presented for the duration of the mouse’s triggering of the center port, and the duration of the center poke required to trigger the start of the trial was gradually increased to 200ms. Delivery of water at the side port was delayed up to 1s with gradual increases as mice became proficient at this training stage. The 3s inter-trial interval was also introduced to decrease repetitive behavior. Mice progressed to the next training stage when they achieved >50% complete trial initiations (remaining for the full 200ms of center port odor presentation) and had rapid reaction times (<2s).
2. **Uncovered Side Training (fig. S1B)**. This stage was not always included but was implemented for animals that had not fully learned the association between center port odors and the required forced side location. Mice completed blocks of ∼20-50 trials directed to the left or right side by the odor provided in the center port with the incorrect side uncovered and accessible. They only received water if they chose the correct side port.
3. **Delay Training (fig. S1B)**. Mice next performed trials with all ports uncovered and pseudorandomly alternating forced left and right trials with the center port odor directing them to the left or the right. All correctly chosen trials were rewarded (1-2 4uL drops) equally on the two sides. The delay between the completion of trial initiation at the center port and water reward delivery following correct choice of the side port was gradually increased by 200-1000ms approximately every 50 trials until the full delay value of 10s was reached. Delays were increased manually while observing mouse behavior to moderate task difficulty and ensure animals maintained adequate motivation. During this stage, a reaction time requirement was instituted such that mice must choose the correct port within a certain amount of time to receive reward. This time was gradually decreased to the final value of 1.2s. Mice moved on to the next training stage when they achieved >70% correct performance at the full delay value.
4. **Introduction of information (fig. S1C)**. In the next stage, trial rewards became probabilistic and side port odors were introduced. Reward probably was decreased to 50% on all trials. 200ms presentations of the side port odors (A,B,C, and D) occurred 1.2s after the go cue in the correctly chosen side port with the appropriate contingencies, such that odor A was always followed by water reward, odor B was never rewarded, and odors C and D were each followed by water reward on 50% of trials. This allowed mice to begin learning that A and B resolved the trial’s outcome and provided information, while C and D did not provide information. 2-3 sessions were performed at 50% reward and then the reward probability was dropped to 25%. Licking responses and the mouse’s presence in the reward port were monitored to ensure they were learning the meanings of the side port odors (mice tended to leave the side port following receipt of odor B). This training stage lasted 5-7 sessions to ensure all mice learned that information was provided.
5. **Choice Training (fig. S1D)**. Mice were explicitly taught that the Choice center port odor allowed them to receive equal probability water reward at either side port in this stage of training. Task event times, side port odors, and reward probabilities were the same as in the previous stage. Mice performed blocks of 10-50 trials with one side port covered to prevent accessing it. Trials with the appropriate Forced odor (Information or No Information) for the uncovered side were alternated pseudorandomly with trials in which the new Choice odor was presented at the center port. Blocks were alternated to ensure mice experienced equal reward rates and total reward amounts following the Choice odor on the left and right side to minimize side bias. This stage lasted for 5-7 sessions, or ∼250-300 Choice trials on each side.
6. **Full Task/Preference Measurement (Fig. 1A)**. Following choice training, mice were tested for preference for information on the full version of the task. Both side ports were uncovered, and mice performed all three trial types, Forced Information, Forced No Information, and Choice, pseudorandomly interleaved throughout each session. Trial types were set in blocks of 12 to ensure mice performed roughly equal numbers of trials of each type within a session, and rewards were assigned in blocks of 8 since trials were rewarded with 25% probability to ensure mice were rewarded frequently enough to maintain their motivation to engage in the task. In initial behavioral tests (n=14), preference was tested for 3 sessions, then the side identities were reversed. Some mice (n=7) performed two additional side reversals such that their preference was tested with information twice on each side for 3 sessions. In mice that were imaged and subsequent mice (n=15), preference was tested for 6 days on each side to ensure stabilized learning of the relevant neural representations. On side reversals, the side locations of the information and no information side port odors were switched while center port odors still indicated movement to the same side. This meant that the center port odor that had previously signaled Information on the left side and was followed by odor A or odor B now signaled No Information on the left side and was followed by odor C or odor D. Similarly, mice had to learn that Information had moved from the right to the left side or vice versa.

#### Water and Delay Titration Experiments (Fig.1, K-N)

Following side reversals, Information was returned to the initial side port and mice were trained for several more sessions (∼3) until their preference re-stabilized. Then a stairstep procedure was used to determine the willingness of mice to sacrifice water reward for information. The reward probability on both sides was raised to 50%, but the reward amount on the information side was changed for 3 sessions at a time. Reward values of 4-ul drops were 1 drop, 6 drops, 2 drops, 5 drops, 3 drops, 4 drops, and then these amounts were re-tested in reverse order. Preference was calculated as the mean across the final session in each block at a given reward amount, such that two sessions were used to calculate the preference at each reward amount. Sessions with 6 drops were omitted from plotting and modeling due to ceiling effects. A similar procedure, but for single blocks of 6 days at each value, was used for imaged mice.

Information preference was measured across different durations of the delay between side odor presentation and water reward using a similar procedure in which preference was tested for 6 days at each delay value: 1s, 10s, 4s, 10s, 6s. Preference was determined as the mean preference for information in the last two sessions at each delay value.

#### Task with Reward Cues on All Trials (fig. S2)

The task was modified to explicitly cue mice when reward would be revealed so that they learned they could leave the side port on all trials. In all stages of training, after mice experienced 80% of the delay length between the side port odor delivery (or in earlier stages, their entry into the side port) and the reward outcome, the correctly chosen reward port illuminated and a tone played for 50ms to indicate whether that trial would be rewarded. This then gave mice 2s at full delay to return to the reward port and collect water. A 500Hz tone indicated a rewarded trial, while a 2000Hz tone indicated an unrewarded trial.

#### Task with Four Information- and Water-Predicting Odors at the Center Port (Fig. 4D)

Two mice with completed surgery for OFC imaging that had been previously trained on the tones task (above) and had strong information preference in that task were trained on this additional task. First, mice were trained to follow the “large water amount” odor to one side port and receive 8 4uL drops of water and the “small water amount” odor to the other side to receive 2 4uL drops of water. The side that had previously been the preferred information side was assigned as the small water side. One side port was covered and mice performed blocks of trials on one side, then the other. Then these two large and small water trial types were pseudorandomly interleaved with all ports uncovered. Due to limitations in the olfactometer, which could only provide 4 odors at the center port, the odor that had previously indicated free choice trials became the “small water” odor, and the fourth odor that had not previously been used was the “large water” odor. Mice performed 7-10 training sessions in this stage. Mice then performed 1-3 sessions of interleaved forced information and forced no information trials as they had learned earlier in the main task version. Mice then performed the full version of this task with information, no information, large water, and small water trials pseudorandomly interleaved in blocks of 8 to ensure adequate exposure to all trial types. Mice performed 6 sessions with the sides in the initial configuration followed by 6 sessions with the side identities reversed. Information and small water were assigned to the same side port, while no information and large water were assigned to the other.

#### Licks

Licks were recorded in initial behavioral sessions (N=14 mice) using capacitance sensors (Phidgets, 1129_1B) at each port’s lick spout.

#### Movement Tracking

Overhead video recording using a machine vision camera (Teledyne, Flea3) was synchronized with behavior using a 10ms digital pulse signal sent to either the Bpod system or Inscopix nVista3.0 imaging system. Video frames were acquired at 20Hz. The mouse’s body position was then tracked using DeepLabCut (*70*).

#### Stereotactic Surgery

Mice were anesthetized with ketamine (100 mg/kg) and xylazine (10mg/kg) through intraperitoneal injection and received analgesia via buprenorphine SR subcutaneous injection (0.75mg/kg) and carprofen (3mg/kg), had fur shaved from their head, and then were placed in a stereotactic frame. Body temperature was maintained using a heating pad attached to a temperature controller. For lens implantation experiments, a 1.1-1.5mm round craniotomy centered on the implantation coordinates was made using a dental drill. Dura was removed and <0.1mm of tissue aspirated, and 0.3uL GCaMP6f virus (AAV1.CaMK2a.GCaMP6f.WPRE. bGHpA, 3.35×10^12 vG/mL, Inscopix) was injected into lOFC (ML: 1.0; AP: 2.4; DV: 2.45mm from Bregma) at ∼50nL/min using a pulled micropipette. After allowing the virus to diffuse undisturbed for 5min, the needle was removed and a 0.5mm or 1mm diameter and 4.0 mm length microendoscope GRIN lens with integrated baseplate (Inscopix) was then inserted at a depth just above the injection site and centered over it (ML: 1.0; AP: 2.4; DV: 2.4mm from Bregma). The combined lens and microscope baseplate assembly (Inscopix) was secured to the skull using Metabond (Parkell) and further secured and covered with black dental cement (Ortho Jet). Mice recovered for at least 1 week before the beginning of water restriction and behavioral training.

#### Histology

Mice were euthanized after anesthesia with ketamine/xylazine by perfusion with 4% paraformaldehyde. Brain tissue was removed for 24hr fixation, and coronal sections (120 um) were cut on a vibratome (Leica). The sections were incubated with far-red neurotrace (640/660, Thermo Fisher Scienti?c) to label neuronal cell bodies. Images were collected using a Zeiss LSM-710 confocal microscope system. Histology was performed to con?rm locations of implanted lenses, as well as expression levels for GCaMP using native fluorescence.

#### Imaging Experiments

Seven mice were imaged throughout the course of training and testing in the main task, including during 10s and 1s delay sessions (tables S1-S5). Two additional mice were imaged throughout learning and performance of the four-odor center port information and water prediction task (JB483, N=198 cells, and JB484, N=239 cells). Recordings of GCaMP activity were made using Inscopix miniaturized microscopes (nVista3.0) with commutators used in passive mode, with uncompressed videos saved using the Inscopix system. Prior to being placed in the behavior chamber, awake mice had the miniature microscope (Inscopix) attached securely to their skull baseplate. Neural activity was then recorded throughout the behavior session (50-100min), and the microscope was removed and the baseplate cover reattached. Images were recorded with the lowest possible LED illumination to visualize calcium transients with settings largely preserved from session-to-session within each animal. Generally, LED power was set between 0.3-1, with most animals remaining <0.7. The Z-focus of the microscope was adjusted rarely to maintain the same field of view based on landmarks such as blood vessels.

#### Quantification and Statistical Analysis

In general, statistical comparisons are marked as ***p<0.001, **p<0.01, and *p<0.05 and the statistical test used indicated in the text. Nonparametric tests were used unless otherwise described. Tests are two-tailed unless otherwise noted.

#### Behavior analysis

Behavior was analyzed using custom Matlab scripts. All mice that completed training during water restriction with adequate weight maintained and that displayed the expected differences in reward port occupancy based on the predicted water reward were included in preference testing. Less than 15% of mice failed to learn the task by this criterion. Mice performed approximately 150 trials per session.

We examined information preference on free choice trials across all of the preference testing sessions (“overall preference”, Figure 1D) or in the last 200 free choice trials prior to the first side reversal and the last 200 trials after reversal before either ending preference testing, reversing the sides again, or progressing to delay or reward amount manipulations. For most animals, these 200 trials occurred in the last three days before and after reversal, since mice performed around 50-60 choice trials per session. Figure S1I shows pre-reversal preference in these 200 choice trials, Figure 1E shows preference before and after reversal. We also examined choice preferences across individual sessions, as in Figure 1C, L, and N. Since choice preference is the probability of the binary outcome of choosing information or no information, we used Matlab’s binofit function to estimate the information choice probability, P(info), and 95% confidence intervals for each animal. We then tested whether the means of these values across animals were different from 50% (indifference) using sign rank tests (Matlab signrank).

We also estimated the probability of correct choice on forced trials using binofit and report data across animals in the same way as preference. For reaction times, reward rate, licks, and probability of poking in nose ports, we report the mean across sessions per animal and computed the population mean and standard error. Reaction time is the time elapsed from the go cue until the first entry of the correctly chosen side port. Reward rate is calculated as the mean water amount received per minute of correct trials. The probability in port is shown as either the mean across the delay between the end of the side port odor stimulus and the reward outcome or the mean across the first 1s after the end of the side port odor stimulus. On plots showing the full trial duration (Fig. 1J and fig. S2G), the trial is binned into 50ms increments to compute the probability of being in a particular port. Distance traveled is computed from the position of the mouse’s nose in each video frame determined using DeepLabCut.

Unless otherwise indicated, behavioral data are from the last three sessions prior to the first side reversal. Prior to pooling data across sessions, we confirmed that there were not significant cross-session differences using a nonparametric within-subjects design (Friedman’s ANOVA with tukey-kramer correction for multiple comparison).

#### Behavioral Models

Cognitive decision models, fit to single trial level-choice data, have been used in neuroscience and behavioral economics for decades to provide evidence that a specific type of computation may be underlying choice trends seen in a given dataset (*71*). The two main questions we sought to answer with decision model frameworks here were (1) “why do information-preferring mice not always choose information?”, and (2) “can the observed experimental behavioral trends be explained by a model that uses separate but interacting reinforcement learning (RL) machinery for processing traditional water rewards and intrinsic information rewards?”. We hypothesized that there is an interaction between RL-like computations of information value and water value following recent literature(*21*) and used a list of relevant models of increasing complexity to test this hypothesis.

In decision model literature, there are a large variety of RL-related models, often becoming quite complex which risks over-parametrization and over-fitting if the number of agents fit and task complexity do not scale with the model complexity (*72, 73*). Thus, for the purposes and constraints of this study, we chose to use the simplest implementation of RL computations, the Rescorla-Wagner model, following similar recent related work in human models (*40*). The models tested are outlined below and incrementally built up to the full model that contains the minimal necessary components to explain both the water-tuning and delay-tuning experiments (Fig. 1, H and I and fig. S2). The MATLAB package fitPsycheCurveWH was used to fit a psychometric curve to each mouse’s real choices as well as the predicted choices from each of the models tested, using the best parameter initializations from a grid search.

First, all models tested transform a decision variable *DV* term, which summarizes the core computation, into a sigmoid-transformed binary choice probability *P*(*t*)_*info*_, here the probability of choosing information at trial *t*:

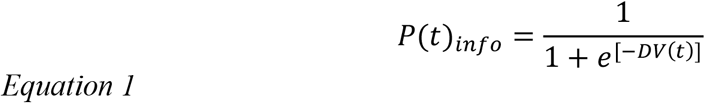

The term *P*(*t*)_*info*_is optimized through a negative

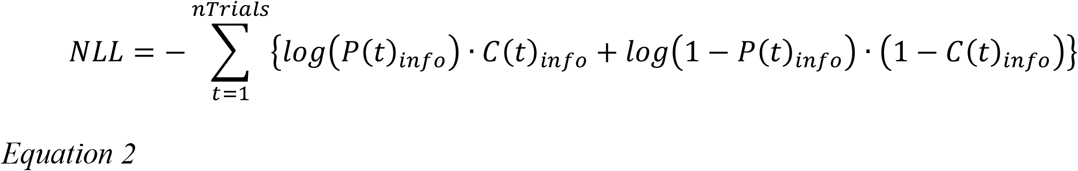

Where is *C*(*t*)_*info*_= 1 for an info choice and = 0 for a non-info choice. Models were each run with 16 initializations of different starting parameter values, and numerically solved using the *fminsearchbnd* in Matlab using a grid search of parameter windows. The nature of *DV(t*) changes in each model, as summarized below:

*RLwater Model:*

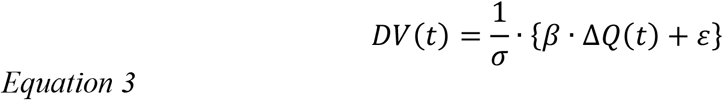

The time-independent fit parameters are *σ*, the normalization constant (inverse temperature), *γ*, a relative weight factor for the traditional Rescorla-Wagner RL term of water Δ*Q*(*t*), and *ε*, a bias term, which allows for a flat baseline tendency towards or away from choosing information that the water term is weighted against. The time-dependent term Δ*Q*(*t*) is defined as the difference of the value functions for the info and no-info sides of the task as usual in binary choice tasks:

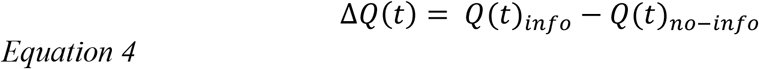

Each *Q*(*t*)_*i*_for *i=info, no-info* is defined according to the Rescorla-Wagner equation with learning rate *α*_1_and prediction error *δ*(*t*) when choice *i* is chosen

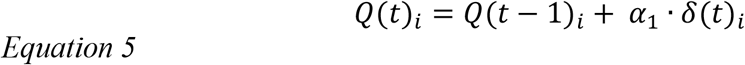

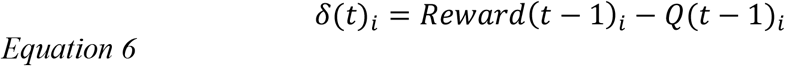

When a choice is not chosen, with unchosen choice *j*, then *Q*(*t*)_*j*_= *Q*(*t* − 1)_*j*_.

The next model tested contains only information-related terms with a new Rescorla-Wagner term *S*(*t*) for information value with information prediction error *θ*(*t*) and learning rate *α*_2_, following analogous logic to recent non-traditional RL equations such as choice prediction errors(*74*):

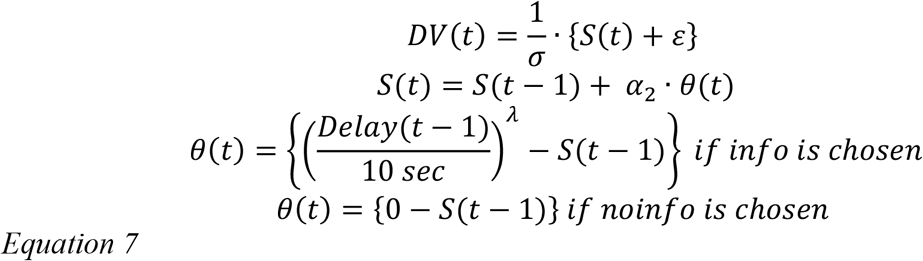

Thus, information’s “value” is 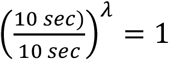 for the maximum delay of 10 seconds, but then is exponentially discounted as delay decreases, following the strong tendency observed in the mice data. When the mice do not choose information, they receive no information value and thus the info-reward is set to zero.

The next model tries to remove the *λ* decay exponent, but combine the *RLwater* term with the *RLinfo* term simplified to have *λ=0*, so that information value = 1 on information choices regardless of delay length.

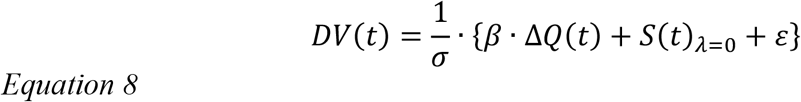

Then, the full model combining both water value and information value computations is defined below, conceptually motivated by recent work finding both types of value are processed in an interacting way(*21*).

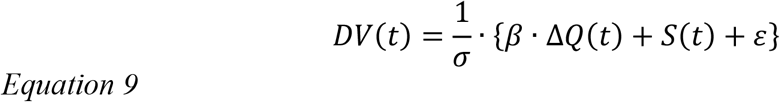

Last, as a conceptually alternative decision model, we made a weighted win-stay lose-shift (WS-LS) model, that regressed the past trial information and reward conditions to predict the current choice:

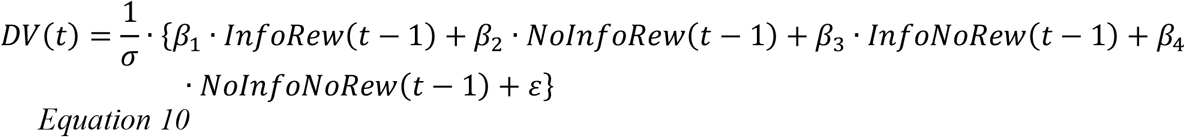

Each *γ*_*i*_condition variable is either 0 or 1. For example, *InfoNoRew* = 1 if the past trial was an unrewarded information choice (either forced or choice), but = 0 else.

The total number of mice fit was *N=*10 for all models, but for 4/10 mice only the water tuning data was fit, not the delay tuning data since those mice did not perform delay-tuned task variants. These mice have had their *λ* discount exponent removed as shown in the Supplementary Table 9 of fit parameters. Then 3/10 mice did not have water tuning data fit. These mice are marked in Supplementary Table 9, but no fit parameters were omitted. To show the data were not overfit, each model was fit five times with different randomized 67% train versus 33% test splits per model for cross-validation (table S11). Mean and standard deviation of normalized AIC and accuracy are shown in the table across the five runs.

To see how much each RL component improved the full model fit, the average Akaike information criterion (AIC) was calculated according to the standard equation for each model with *k* as the number of free parameters(*75*):

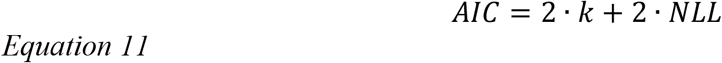

The full *RLwater + λ-RLinfo* model is presented in Figure 1. The auxiliary models are presented in figure S3. It is visually clear that only the *RLwater + λ-RLinfo* can simultaneously capture the important trends from the data in both water tuning and delay tuning experiments. Future work seeking to do more advanced decision modeling with further corrections to the Rescorla-Wagner formulation should increase the sample size and tune a wider range of experimental conditions, but the current models implemented show that the minimally simple RL formulation that accounts for both water value and information value is able to explain the full range of experimental conditions tested here. As a caveat, we do not claim our fitted parameters are unique or will work for all future studies, a common issue in cognitive decision models because model parameter magnitudes and scaling are not independent and thus degenerate solutions likely exist (*71, 72*). However, this well-known issue does not undermine the finding that the minimal two-value-type RL model machinery performs well in our context.

#### Imaging Data Processing

Raw video data were spatially downsampled at 4x and saved as Tiff files using the Inscopix Python API. We used the NormCorre rigid motion algorithm (*76*) to both remove movement artifacts from imaging videos and to align the field of view across sessions to facilitate cell registration. We motion corrected videos filtered to reveal more stationary landmarks such as blood vessels and then applied the computed X-Y shifts to the original videos. To filter videos, we took a difference of Gaussians approach to extract potential stationary features smaller than cell diameter from the larger background fluctuation. We applied a Gaussian filter (Matlab imgaussfilt) with radius 2 to our downsampled 200×320 pixel images and subtracted pixel values of the images separately filtered with a larger radius (Matlab imgaussfilt radius 6). We then set a threshold value of 4 and set all pixel values less than this threshold to its value. This resulted in a filtered video with mostly black background and light-to-white landmarks on which we performed motion correction and generated a mean template image.

Motion correction was performed within each session using the NormCorre algorithm using the following parameters: ‘bin_width’, 500, ‘init_batch’, 1000, ’max_shift’,10, ‘iter’,2. We then used NormCorre to align each session’s video frame-by-frame to the filtered template from a central “anchor” session for each animal (typically the final day of preference testing before the first side reversal). The alignment of the template field of view from each session was manually inspected to ensure accurate registration across sessions. Cell location footprints and activity were then extracted using the Matlab implementation of CNMF-E (*77*) as follows.

CNMF-E default parameters were used, except that min_corr and min_pnr were adjusted for each animal to maximize cells identified during the initialized step of the algorithm (min_corr range 0.7-0.8, min_pnr 8-10) and gSig=3, gSiz=11. For initialization, min_pixel was set to 25 and for identification of cells in the residual after background subtraction, min_corr_res = 0.8 and min_pnr_res = 9. Following two iterations of CNMF-E’s cell footprint and calcium activity extraction, we applied several additional filters to remove false positives, taken from the incorporation of the CNMF-E algorithm into the Caiman Python implementation (*78*). We set thresholds for temporal sparseness, minimum peak-to-noise, and a constant baseline of activity. We first removed cells with temporal sparsity < 0.003.

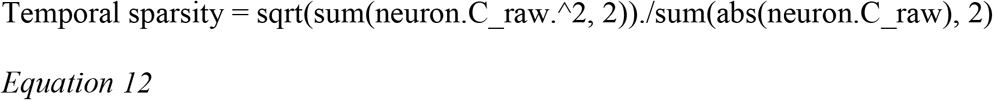

We then removed cells with peak_to_noise_ratio < 8.5

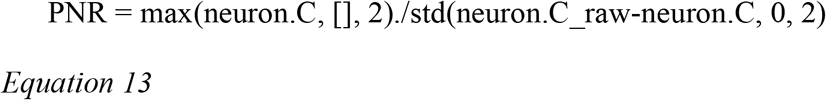

Finally, we removed cells in which CNMF-E’s detrending of the calcium fluorescence across the session was ineffective and that had large differences in the baseline fluorescence between the beginning and end of the session. We computed the difference from baseline by taking the difference in mean activity between the first and last 10% of the session frames and removed cells for which this absolute value was greater than 3.

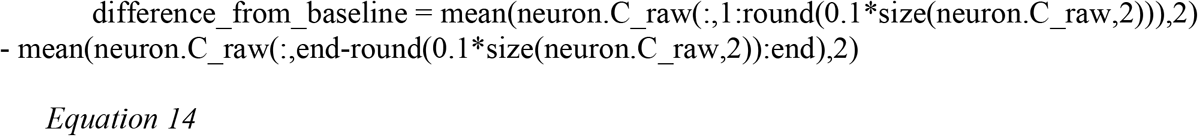

We then visually inspected each cell using CNMF-E’s viewNeurons method to ensure appropriate neuron-like shape and calcium transients and removed false positive cell identifications.

Using this procedure, we were able to register cell spatial footprints across sessions several weeks apart, over the entire months-long course of training, for up to 4 sessions simultaneously using the register_multisession function from the Matlab implementation of CaImAn. (*78*). We used the default parameters for register_multisession, with the exception of setting the threshold for turning spatial components into binary masks, options.dist_maxthr=0.22, and the threshold for setting a distance to infinity, options.dist_thr=0.6. Cell registration fidelity between individual sessions was visually inspected using the register_ROIS method for maximum overlap and false positives to appropriately set parameters. For mice imaged in the 4 center port odor information and water value prediction task, a newer algorithm, SCOUT, was used to register cells since only two animals were imaged. SCOUT registration of cells used the default parameters (*79*).

Neural activity in each session was aligned to behavioral events using DAQ timestamps, and we analyzed neural activity across all mice considering cells across animals as a single pool for single-cell analyses. Our CEBRA model simultaneously fit across animals using all cells, not just those registered day-to-day, with the same qualitative findings about information and water value encoding supports pooling cells across animals. The main dataset (table S1) consisted of cells registered across 4 sessions surrounding a side reversal such that there were 2 sessions with information on each side (left and right), with 56-216 cells per animal across 7 animals (N=1138 total neurons). Datasets for delay comparisons (10s vs. 1s, tables S3-5)) and learning (table S2) also consisted of 4 sessions per animal across 6 and 7 animals, respectively, with similar numbers of cells per animal (N=992 total delay neurons, 788 total learning neurons). Data for mice imaged in the four odor center port information and water value prediction task involved cells registered across 4 sessions surrounding a left-right side reversal, with 437 cells.

#### Analysis of Neural Activity

We used the calcium signal, C, output by CNMF-E that is smoothed with a kernel based on the time course of GCaMP6f signal decay as the calcium activity, a proxy reflecting neural spiking activity, of each cell across each session. Because CNMF-E subtracts the background signal from each imaged frame individually, the output activity, C, may be considered a scaled version of the change in fluorescence over background (ΔF/F) at each frame. Moreover, given this background subtraction, this signal is already normalized (*77*). We therefore analyzed and report raw “calcium activity.” We also tested a standard z-score normalization procedure subtracting each cell’s mean across each session and dividing by the standard deviation, and this did not qualitatively affect the results.

We computed indices of cross-validated differential activity to analyze OFC neurons’ encoding of task variables (coding index). For each pair of conditions (e.g. information and no information), we balanced (cross-validated) the direction of the difference in activity (i.e. sign). We multiplied the mean difference in activity between the two conditions for one arbitrary half of the trials (e.g. even numbered trials) by the sign of the mean difference on the other half of the trials (e.g. odd numbered trials) and vice versa and then took the mean value of these two split halves as the coding index.

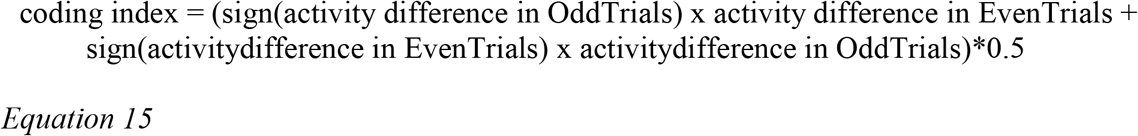

This procedure is similar to taking the absolute value of the difference in activity but instead creates an index of differential activity centered at zero in the absence of consistently greater activity in a particular condition.

We computed the coding indices across the full trial time course at each imaged frame, beginning at the onset of center port odor on the mouse’s center port entry that successfully initiated the trial. Mice sometimes entered the center port and received a partial odor stimulus prior to successfully initiating a trial by remaining in the port through the completion of the go cue tone, which may have meant that OFC responded differently at the animal’s first and subsequent center port odor presentations. Thus, we computed full-trial coding indices surrounding the odor presentation at successful trial initiation, from which the remaining trial events proceeded.

To analyze neural responses at specific trial epochs, we examined a 1-second window following each trial event: the first center port odor presentation, the side port odor presentation, and the reward outcome. We determined the 1-second post-event activity window by observing the raw odor responses and peak differential activity around each event in our task. We also determined the time course of odor stimulus delivery in the nosepoke ports using a photoionization detector (Aurora Scientific, 200b miniPID). Although we observed small differences in the timing of odor availability, on average odor was first detectable 0.075s after odor valves were open and peaked at 250ms after valve opening. The neural responses we observed peaked within 1s of odor onset. We have therefore used a window of 0.2-1.2s after the timing of events recorded by our Bpod system (odor and water valve opening), and a comparable window 1.2-0.2s before events, to analyze neural responses. For plot visualizations, we have indicated odor onsets 0.075s after the opening of the odor valves in the olfactometer.

To determine whether cells, and the OFC population on average, encoded differential activity between two conditions as computed in our coding index in a task epoch, we took a bootstrapping approach. We compared the difference in the mean coding index in the 1s window after an event to the mean index in the 1s window before it to account for the fact that activity continuously varied throughout the task and in particular, was often already elevated just prior to the side port odor delivery. Thus, for a given set of conditions (e.g. information and no information), we shuffled trials between the two conditions 1000 times. For each shuffle, we determined the difference between the mean coding index in the 1s window after the trial event and the mean coding index in the 1s window before the trial event. We then determined the percentage of these shuffled values that were less than the observed increase in differential activity to estimate the statistical likelihood of the observed difference (p-value). We defined a cell as contributing to a representation or encoding a difference between conditions if the observed coding index difference was greater than that of at least 5% of shuffled data. Similarly, we defined that the OFC population encodes a difference between conditions if the mean population coding index difference around an event was greater than 95% of the shuffled values. Figure S4D-G shows the event-specific coding indices we computed in addition to the full trial information and water coding indices.

#### Cells with significant conditional responses to task events (responding cells)

Cells were identified as responding to an event in a task condition using a rank sum test to determine if their activity in the post-event 1s period (as above) in that condition was different from activity in the pre-event 1s period in that condition, which we considered the baseline activity for that event. To accommodate baseline activity that was already elevated due to earlier trial events given the close proximity in time of the center and side port odors, we determined the maximum activity for each cell on each trial in the pre-event period and fit an exponential decay function based on GCaMP6f fluorescence (λ=4) to determine its predicted activity in the post-stimulus period. If the absolute value of difference between the mean of that predicted activity in the post-event period and the mean of the observed activity in the post-event period on that trial was greater than 0.2, the GCaMP fluorescence-predicted activity in the post-event period was used as the baseline. If not, the pre-event activity served as the baseline. Cells with significant responses were then identified as those in that condition with mean activity in the post-event period significantly greater than baseline using a rank-sum test and with the value of that difference greater than 0.1. This procedure offers a conservative identification of responding cells in order to prevent false positives. Using a more standard procedure with a rank sum test on mean activity before and after the event falsely identified many cells with delayed responses to the center port odor whose activity then diminished as expected for calcium signals at the same time as the side port odor delivery as “responding” to the side port odor. We therefore used the above approach to more conservatively identify cells with activity changes due to the side port odor.

#### Mean activity

For plots showing population mean activity, each cell’s mean activity on the indicated trial type was calculated, its mean activity within the 1-second pre-event window was subtracted, and then all cells’ mean activities were averaged. Single cell plots are shown with the mean activity in the 1-second pre-event window subtracted. To test whether the population mean responses to center port odors changed across delay changes and learning (Figure S6A,K), we compared the mean activity in the 1s window after the center port odor across all trials between the two conditions, using a rank sum test for significance.

#### Heatmaps

Population calcium activity and coding index heatmap plots display the mean activity or coding index for each cell across trials in each condition, with the cell’s mean activity/coding index in that condition across the pre-event 1s period subtracted. Difference heatmaps show differences in mean activity or coding index between conditions without pre-event mean subtraction. Plots within panels are all sorted by the indicated activity such that a single cell can be read across the entire heatmap.

#### Decoding

Linear classifiers (support vector machines) were used to decode trial types. Classifiers (Matlab function fitclinear, default support vector machine) were trained over 100 iterations on 80% of data and then tested on 20% of held out data. Trials were balanced between relevant types. Classification was performed on 200ms bins of neural data, with calcium activity averaged for each cell on each trial within each bin. Because trials were pseudorandomized with a rigid block structure and mice had to repeat incorrect trials, decoding performance could be greater than chance prior to the odor stimulus delivery. We therefore determined the statistical significance of the improvement in decoding accuracy following the odor stimulus by comparing the mean accuracy across animals in the 1s interval after odor to the mean accuracy in the 1s interval before odor across animals in the main task using a Wilcoxon sign rank test. For the decoders trained on the 4 center port odor task variant, since data were only available from two animals, we assessed the improvement in accuracy following the center port odor using a Wilcoxon sign rank test to compare the difference in mean accuracy in the 1s window before and after odor across the 100 iterations of classifier training and testing. Information and water decoding cells were determined by taking the absolute value of classifier weights from the classifiers trained on information vs. no information and large vs small water trials, respectively, and identifying cells in the top 10%. Classifiers were then trained using different subsets of cells based on these identities and accuracy determined in the time period 0.8-1s after center port odor onset. For subsets excluding the information or water cells, 10 iterations of a balanced number of randomly selected cells were performed. Significant differences in accuracy were determined using a kruskal-wallis test across 100 iterations of train-test across the different classifiers with a tukey-kramer correction for multiple comparisons.

#### PCA of information-no information axis

To compute an axis along the information-no information dimension in population activity space, we calculated the difference between the mean activity on information forced and the mean on no information forced trials for each cell following the center port odor presentation (0-0.8s after odor valve opening). We then computed the principal components of that population activity difference (Matlab function svd). The first component captured >80% of the variance in this activity computed from the variance of the projection of the information and no information trial activity onto the first component divided by the total variance of activity on information and no information trials. We therefore projected the population activity in each trial, and the mean within each condition (information and no information) onto that component for visualization.

#### CEBRA Modeling

We used Consistent EmBeddings of high-dimensional Recordings using Auxiliary variables (CEBRA) to better understand how a complex task such as ours is represented at the neural level. The complexity of our task, together with the possibility that the OFC, as a frontal region, contains neural representations that are more compressed and processed compared to other regions, suggested that neural task representations are more likely to have arisen from the non-linear combination of neural signals (*33, 36*). A nonlinear latent variable model such as CEBRA, unlike principal components analysis (PCA), for example, reduces the chance that an analysis is fitting task-irrelevant noise or missing important nonlinear signals that are crucial to accurately capture how a task is encoded at the population level.

More specifically, we used CEBRA to obtain low-dimensional task embeddings by jointly fitting neural data and task-relevant behavioral variables in a time-dependent manner. We generated time-stamped labels expressing task structure and mouse behavior, as further specified below, and fitted these together with the neural data by minimizing an InfoNCE contrastive loss objective (*51*). We fit the model to the entire cohort of animals at once, such that contrastive samples were chosen from across all animals. We thus obtained latent embeddings that were informative and consistent across animals. We examined the geometry of the information and water value representations in the embeddings and measured the differences between these two representations over the course of each trial. We confirmed that the embeddings were informative by decoding task-relevant variables and comparing results to shuffled versions which scrambled the association of particular labels with the recordings.

We fit CEBRA models with the following specifications: model architecture = “offset1-model”, batch size = 1024, distance = ‘cosine’, conditional = ‘time_delta’, temperature = ‘auto’, learning rate = 0.001, max iterations = 5000, output dimension = 3, number hidden units = 100. We arrived at these settings by fitting CEBRA models on the data from 3 mice and picking the settings that achieved the lowest overall InfoNCE (*80*) loss value after 10000 iterations; during this process we observed that only half the iterations were necessary for the loss to converge so we set max iterations = 5000 to minimize compute.

We fit two types of models which we refer to as *full* and *delay*. The full model is fit on data with a long delay (t_d_l = 10 seconds) between the side port odor and reward delivery, while the delay model is fit on data with both short (t_d_s = 1 second) and long delay (t_d_l = 10 seconds) and is aimed at embedding the two delay types in a joint space for comparison.

The full model was fit on 6 mice (JB432, JB426, JB434, JB413, JB424, JB425) with cell and trial counts as specified below (table S6). Importantly, we employed a multi-session setup in which each session with its unique dimensionality was added into the fitting process accompanied by shared context labels (table S8) which allowed CEBRA to learn a shared embedding space for signals from all sessions (across all mice). We used sessions from our main data set balanced before and after information side reversal, such that each mouse was fit with two sessions with information on each side (left/right). In order to achieve an unbiased representation across all conditions we equilibrated the number of trials across conditions such that there were equal numbers of trials across the four conditions resulting from the interaction of “reward vs. no reward” and “information vs. no information.” For the full model, the number of time points per trial is fixed at n_time = 320 time points per trial throughout, ranging from the initiation of the last (if multiple) presentation of the center port odor all the way until after reward delivery.

The *delay* model was fit on 6 mice (JB432, JB433, JB434, JB426, JB424, JB425) with cell and trial counts (table S7) and label types (table S8) with n_time=113, as specified below. In order to make the two delay types (*short* and *long* delay) comparable, with identical context labels across sessions allowing sessions with variable cell counts to be fitted together in one model, we excerpted the outcome period in the *long* delay trials and concatenated it after the first ∼1s of the 10s delay period (omitting most of the longer delay) thus making the overall duration of short and long delay trials the same. These sessions do not include a reversal of the information port location. As before, in order to achieve an unbiased representation across all conditions we equilibrated the number of trials across conditions such that there were equal numbers of trials across the four conditions resulting from the interaction of “reward vs. no reward” and “information vs. no information”; in case that resulted in less than 1 trial per condition, we equilibrated instead by “information vs. no information” only (session marked with * in table S7).

Unlike the *full* model, we matched cells across all recording sessions for a particular mouse before fitting the *delay* model (hence the matching cell counts in table S8). We then combined all of a mouse’s recording sessions into a single array and fitted the recordings and labels (table S8) with CEBRA in a *single*-session setup. This allowed us to derive a common embedding space across both delay types.

We fit all CEBRA models on the neural data (as defined above) together with the following label types (table S8). Labels are zero everywhere except between the start and end frame where they are set to nonzero values depending on label type.

We obtained the decoding performance for the CEBRA embeddings. We divided the data from each animal and session separately into a train and a test set such that data from each animal and session was present in both sets, with 40% of the data being assigned to the test set. We obtained a CEBRA fit on the train set with the parameters described above and then projected the train set recordings into the embedding space. We thus obtained 3-dimensional embeddings for every trial in the train set and used these trajectories to train a k-Nearest Neighbors classifier with a cosine-distance metric and k=20. We fitted separate classifiers for each label type to be decoded. We then used these classifiers to predict a given label type at each point in a trial for all the recordings in the test set. To quantify performance, we calculated the balanced accuracy score across a bin of 5 samples with a step size of 1, and averaged results across sessions, animals and runs. The results we report are averaged across 5 runs, meaning 5 separate CEBRA fits with different random seeds. We calculated the 95%-confidence interval for the decoded accuracy results across the six animals, accounting for dependent measurements across sessions (*81*).

We also fit label-level controls. There were 4 different label-level control types: “shuffled trials all labels”, “shuffled max all labels”, “shuffled info label” and “shuffled reward label”. For the “shuffled trials all labels” condition, we randomly scrambled the assignment of trials to their original time-varying labels, for all the label types, but left the relation between labels and the time they were assigned in a trial (table S8) intact. For the “shuffled max all labels” condition, we randomly scrambled both the trial- and the time-affiliation of a given label, for all label types, thus resulting in the maximum amount of signal scrambling. For the “shuffled info label” condition, we only scrambled the trial-label assignment for the “info vs no info” label type (table S8), leaving all others intact. For the “shuffled reward label” condition, we only scrambled the trial-label assignment for the “rewarded vs unrewarded” label (table S8), leaving all others intact. Otherwise, controls were fitted identically to the full model. Results are again reported as an average across 5 runs, using different random seeds to achieve the train/test allocation (fig. S8).

In addition, we fit cell-level controls. There were two different cell-level control types, 60% and 20%, with the percentage number indicating how much of the original population of recorded cells was used (tables S6 and S7). So, for instance, in the 60% condition, only 60% of the cells in a given session were randomly picked to be included in the fit. This was also repeated across 5 runs, using 5 different random seeds to subsample the cells. Results are reported averaged across sessions, animals and runs.

Finally, we obtained the distance between individual trajectories in the embedding space. For that we projected all the recordings into the embedding space obtained by the *full* CEBRA model fit. To determine the distance in neural population space between task embeddings in the various experimental conditions (“info”, “no info”, …), the Euclidean distance was calculated between all possible trial pairs. Cross-session and cross-animal first and second moments were calculated by inverse-variance weighting. Averaged cross-session variances and cross-animal variance were added to obtain the final 95% confidence interval (CI).

**Fig. S1.**
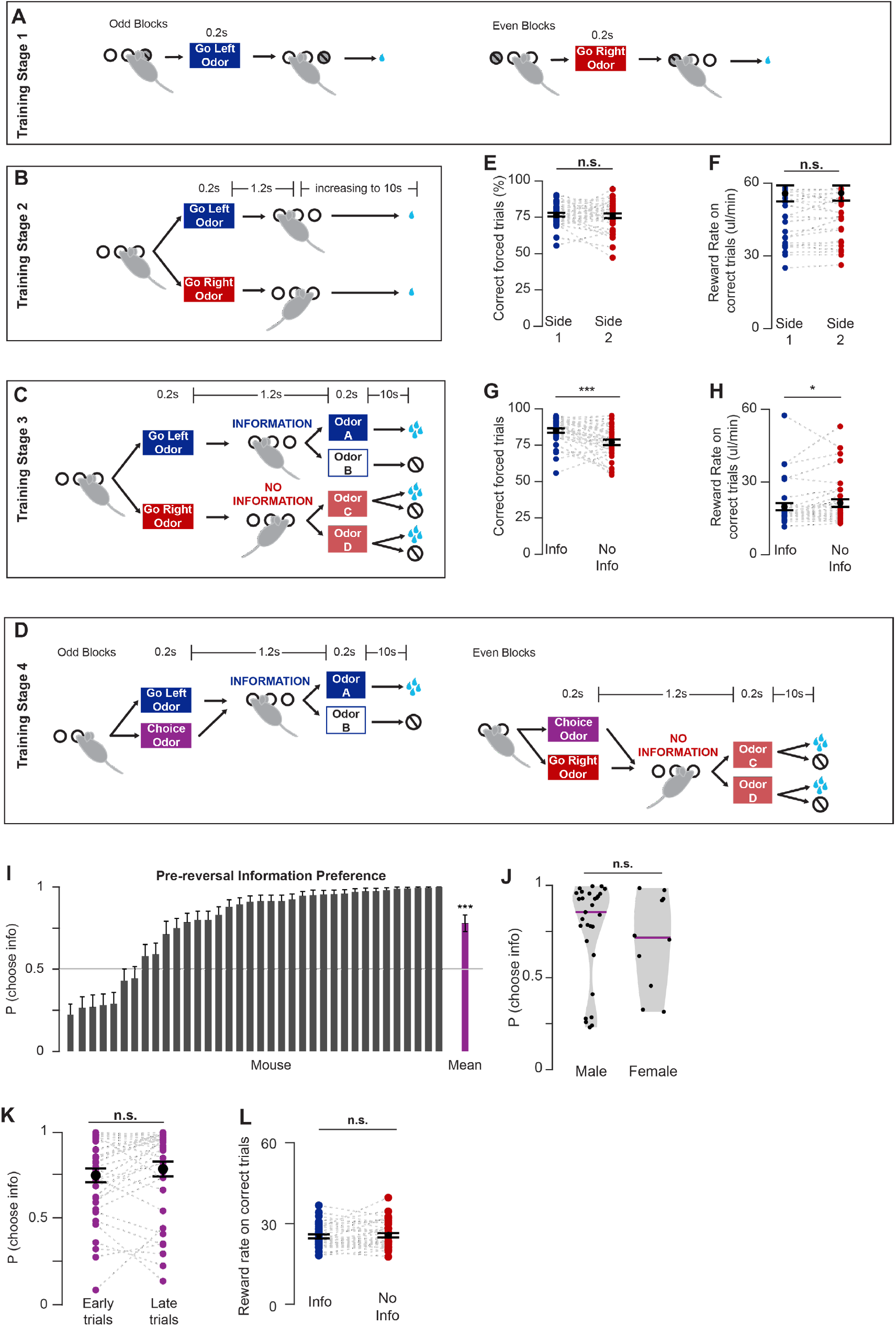
(**A**) Initial task training stage with alternating blocks with one side port covered. Mice learn to first receive odor in the center port and then enter the uncovered side port. (**B**) Second task training stage in which mice learn that odor directs them to either the left or right side on pseudo-alternating trials and the delay to reward are gradually increased. (**C**) Third task training stage in which rewards are made probabilistic and side port odors (information and no information) are introduced. (**D**) Fourth task training stage in which the free choice center port odor is introduced in alternating blocks of trials on the information or no information side such that mice learn they can choose either side on free choice trials. (**E**) Correct performance of forced trials in training stage 2 when both sides offer identical water reward and no information, points for each mouse, black for mean +/-std err across animals, Wilcoxon sign-rank. (**F**) Reward rate on forced side 1 (future information) vs. side 2 (future no information) trials in training stage 2 when both sides offer identical water reward and no information, Wilcoxon sign-rank. (**G**) Correct performance of forced trials in training stage 3 when information is introduced. (**H**) Reward rate on forced information vs. forced no information side trials in training stage 3, Wilcoxon sign-rank. (**I**) Initial information preference. Estimated probability of choosing information (binomial fit) and 95% confidence interval on the last up to 200 free choice trials prior to the reversal of side identities for each mouse. For each mouse, estimated probability of choosing information (binomial fit) and 95% confidence interval is shown. Mean shows mean and std err of estimates across animals, preference different from 0.5, p<0.001, Wilcoxon sign-rank. (**J**) Pre-reversal choice of information (last 3 testing sessions prior to side reversal) for male and female mice. Points for each mouse, purple line indicates median, Wilcoxon sign-rank. (**K**) Choice of information on trials in first vs. last third of session in last 3 testing sessions prior to side reversal. We observed no effect of satiety, Wilcoxon sign-rank. (**L**) Reward rate on correct forced information vs. forced no information trials in last 3 testing sessions prior to side reversal, Wilcoxon sign-rank.

**Fig. S2.**
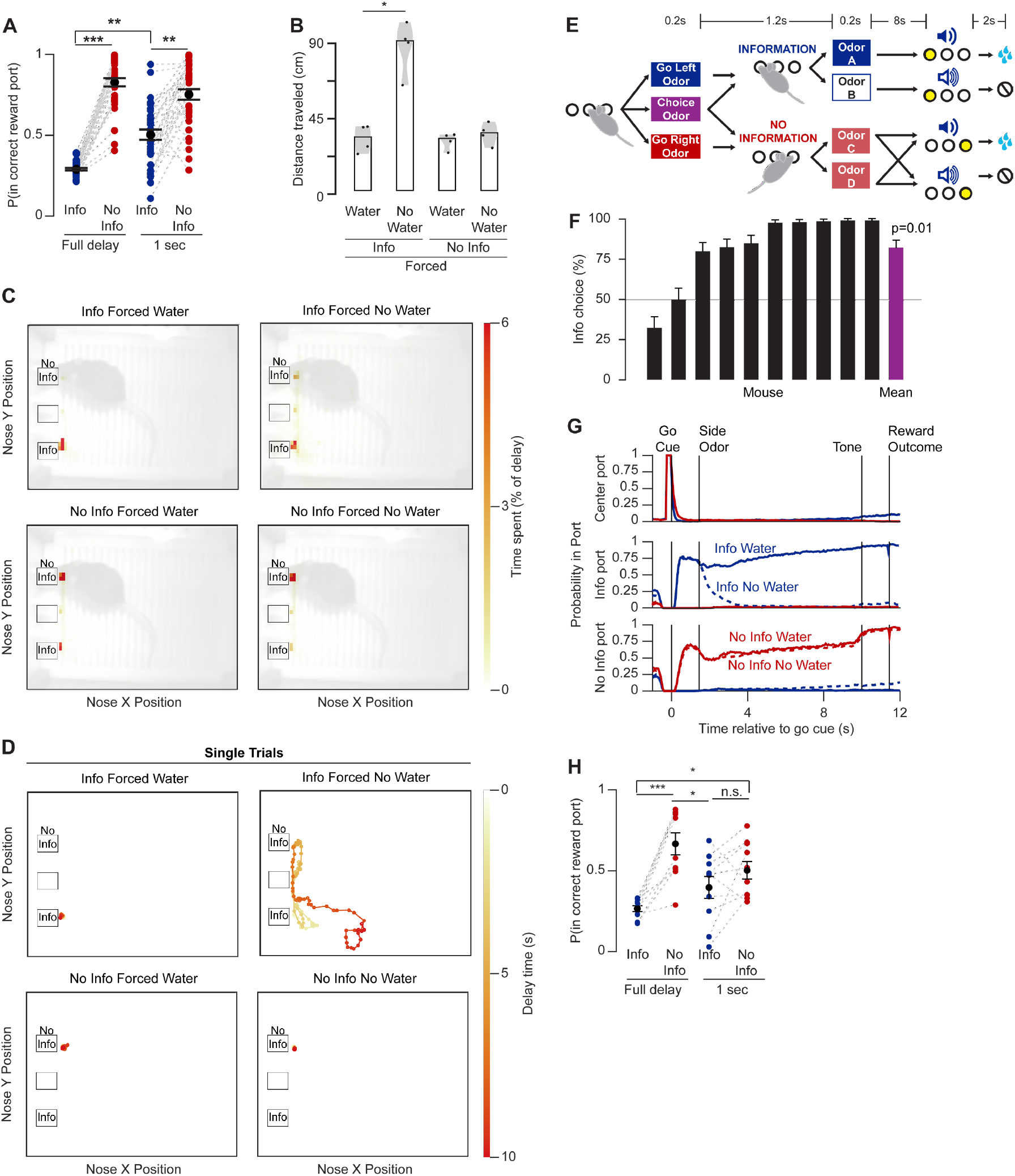
(**A**) Probability of mouse remaining with nose poked in the correctly chosen side port on forced information vs. no information trials, across the full delay from the end of side port odor delivery to reward outcome and in the first second following side port odor delivery, points show the mean for each mouse, black shows mean +/-std err across animals, Friedman’s ANOVA with tukey-cramer correction for multiple comparisons. (**B**) Distance traveled during the delay by trial type for each mouse. Bar shows mean across animals, Friedman’s ANOVA with tukey-cramer correction for multiple comparisons. (**C**) Spatial distribution of mouse’s nose position during the delay between side port choice and reward outcome by trial type for sessions with information on the left side, N=4 mice. Position of nosepoke ports shown as black outlined boxes. (**D**) Representative single-trial trajectories of mouse’s nose position during the delay by trial type. (**E**) Task variant with cue for outcome delivery on all trials. The correct reward port was illuminated and a tone indicated whether the trial was rewarded or unrewarded 2s prior to possible water delivery on all trials. (**F**) Initial information preference in the cued outcome task variant. Choice of information on the last 200 trials prior to side reversal (mean binomial fit and 95% confidence interval shown for each mouse, mean shows mean preference and CI across animals). (**G**) Probability of mouse nose poking in each port by trial type in the cued outcome task variant, N=8. (**H**) Probability of mouse remaining with nose poking in the correctly chosen side port in the cued outcome task variant on forced information vs. no information trials, across the full delay from the end of odor delivery to reward outcome and in the first second following odor delivery. Points for each mouse, black for mean +/-std err across animals, Friedman’s ANOVA with tukey-cramer correction for multiple comparisons.

**Fig. S3.**
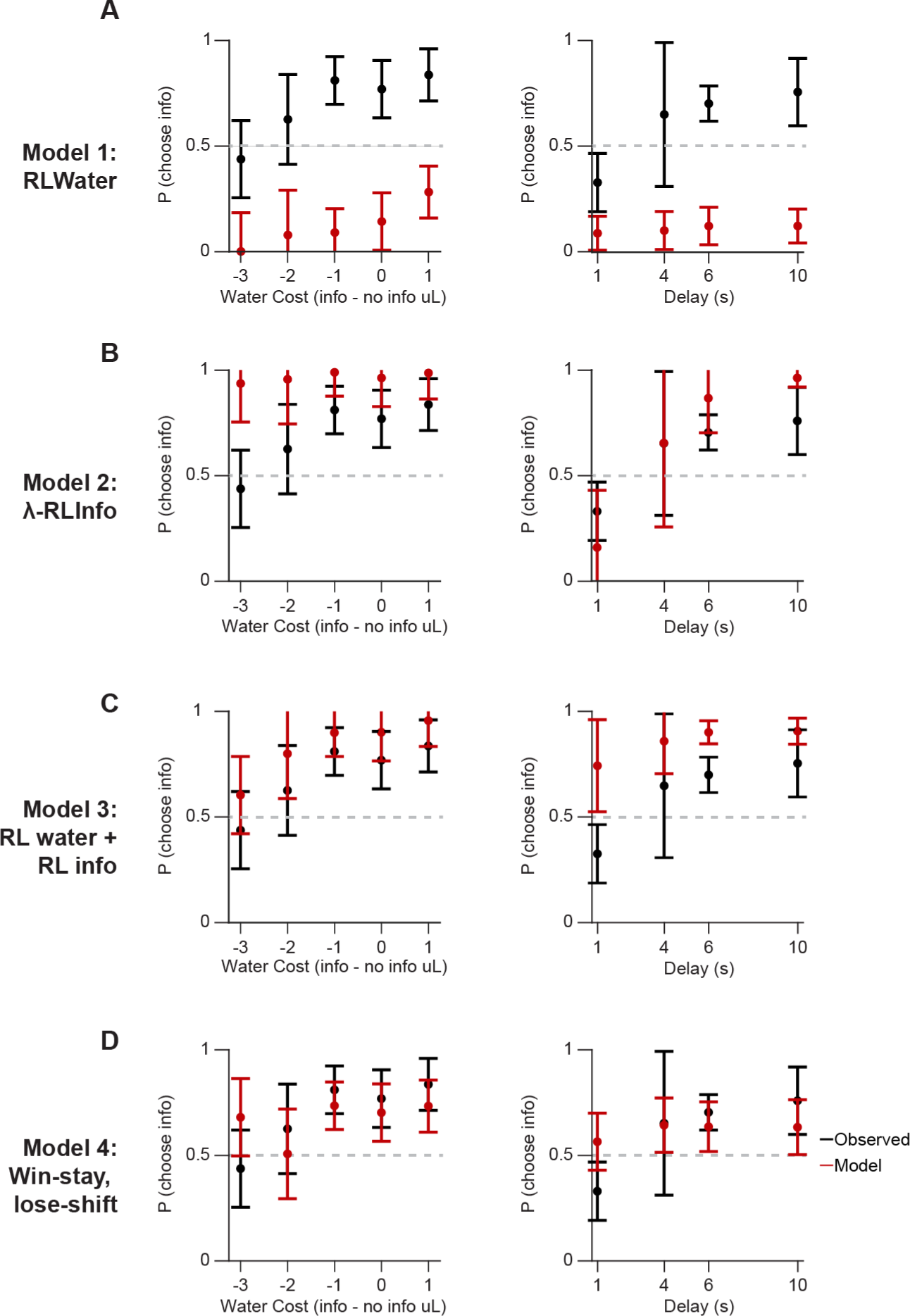
An ablation study of our decision model in Fig. 1 and Methods was performed to show the behavioral effects of dropping each of the key model terms and variables. (**A**) When the information value Rescorla-Wagner term was dropped (RLwater), the ability to predict information preference completely collapsed. The predicted preference dropped below 50% because the model was fit to actual mouse choices and trial variables, and the mice on average preferred information. (**B**) When the Rescorla-Wagner term of water value was dropped but the delay-discounted Information term was kept (λ-RLinfo), the model overestimated the information preference at the highest delay value and underestimated the Information preference at the smallest delay value. It also completely failed to predict the behavior in the water value-tuning experiments. (**C**) When the discount exponent was dropped on the information value term, so that the information value was unchanged at all delay durations (RLwater + RLinfo), the model failed to predict the results of the delay tuning experiment. (**D**) Last, we tested a win-stay, lose-shift model where mice have sigmoid transformed regression coefficients for each possible previous trial type and outcome (info rewarded, no-info rewarded, info unrewarded, no-info unrewarded). For example, mice may want to switch more often after unrewarded trials. This model predicted overall average behavior in a coarse-grained way, but failed to predict behavioral changes as relative water value and delay duration were tuned. Error bars are 95% CI.

**Fig. S4.**
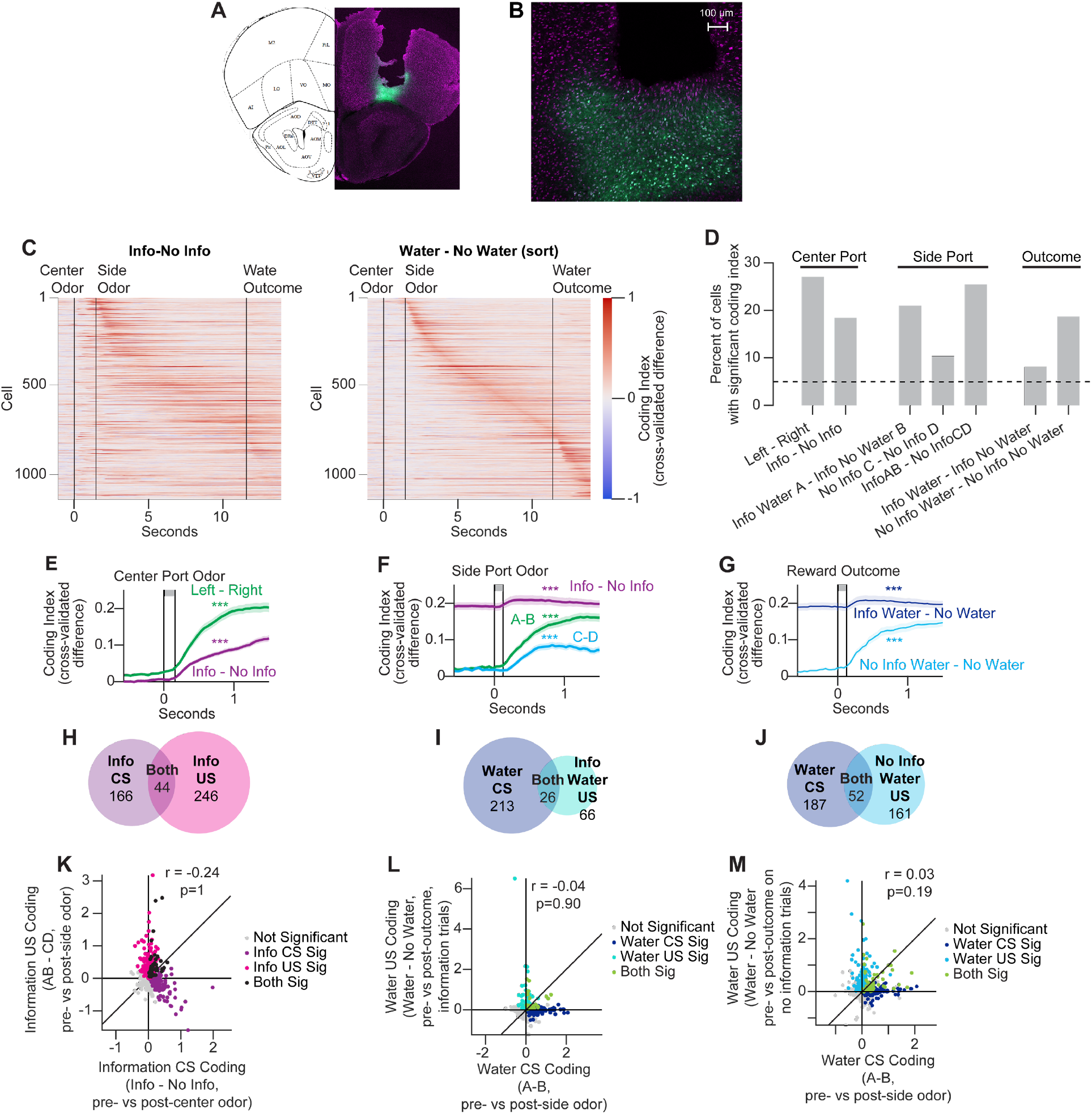
(**A**) Representative image of GCaMP-expressing virus (green) and GRIN lens location, with neuronal cell bodies in magenta. Coronal section of an example brain and aligned atlas view. (**B**) Magnified view of A. (**C**) Left, mean information value coding index across the population. Right, mean water value coding index \. Data as in Figure 2B, except here cells sorted in both by water coding index. (**D**) Percent of cells (N=1138) with significant encoding of each of the indicated types of differential activity. Activity significance measured as the mean cross-validated difference (coding index, see Methods) between the two conditions in 1s interval after the indicated event compared to 1s interval before. Differences compared to shuffled data for significance, p<0.05. (**E-G**) Population mean coding indices. Shaded area shows standard error. (E) Coding at center port odor, (F) at side port odor, (G) at reward outcome. Difference in mean index in 1s interval after the indicated event - 1s interval before compared to shuffled data for significance. C-D activity difference reflects that odor D is provided on 25%, whereas odor C is provided on 75%, of no information trials. (**H-J**) Number of cells with significant coding as in (D) and their overlap, all p>0.05, chi-square test, N=1138. (**H**) Info CS (information – no information at the center port) vs. info US (info trials with odors A and B – no info trials with odors C and D at the side port). (**I**) Water CS (odor A – odor B at the information side port) vs. water US on information trials (water – no water outcome on information trials). (**J**) Water CS (odor A – odor B at the information side port) vs. water US on no information trials (water – no water outcome on no information trials). (**K-M**) Correlation between CS and US coding for (K) information, (L) water on information trials, (M) water on no information trials. Each point represents the mean of the indicated coding index in 1s interval after the indicated event – the mean in 1s interval before. Cells colored by significant coding, Pearson correlation values shown.

**Fig. S5.**
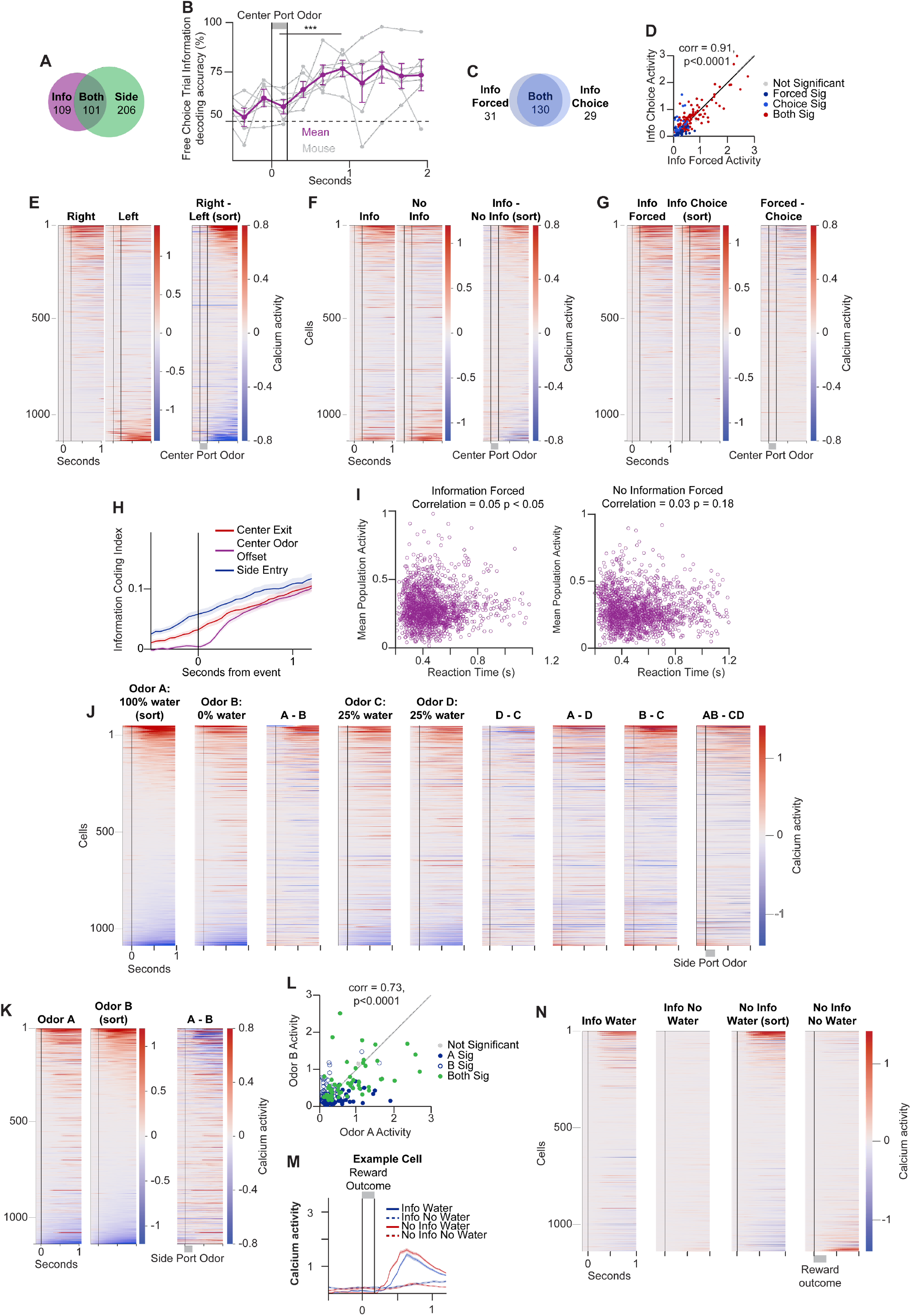
(**A**) Number of cells with significant coding of information and side at the center port and their overlap, all p>0.05, chi-square test, N=1138. Coding computed as the difference in mean index in 1s interval after – 1s interval before the center port odor with significance (p<0.05) determined by comparison with shuffled data. (**B**) Accuracy of decoding of held out free choice information versus free choice no information trials with a linear classifier trained on free choice information and no information trials. Purple, mean +/-95% CI across mice, gray=individual animals. Mean accuracy greater in 1s interval after center port odor than before, Wilcoxon sign-rank. (**C**) Overlap of cells responding to center port odor on forced and free choice information trials (N=1138, p<0.001, chi-square test). Response defined as greater mean activity in 1s following odor than 1s before, see Methods for details. (**D**) Correlation between center port odor responses (mean activity in 1s after – 1s before odor, “baseline”, see Methods) on forced and choice information trials. Pearson’s correlation shown. (**E**) Population responses to center port odors directing to the right or left side port. Each cell’s mean activity on left and right forced trials, and the difference between them, shown. Cells in all columns sorted by difference. (**F**) Population responses to center port odors directing to information or no information on forced trials. Plots as in (E). (**G**) Population responses to center port odors on forced and choice information trials and their difference. Plots as in (**E**) but sorted by information choice trial activity. (**H**) Population information coding activity aligned to choice-related events. Lines show mean information coding index; shading shows standard error. Time 0 corresponds to the indicated event. (**I**) Correlation between OFC activity and reaction time on, left, forced information and, right, forced no information trials. Each point shows the mean activity across all cells in 1s interval after center odor on a single trial. Pearson’s correlation shown. (**J**) Population responses to side port odors and differences between mean activity. Each cell’s mean activity on trials with the indicated odor, or the difference in mean activity between indicated odors, shown. All columns sorted by odor A activity. (**K**) Population responses to side port odors A and B, sorted by response to odor B. (**L**) Correlation between side port odor responses (mean activity in 1s interval after – 1s interval before odor, “baseline”, see Methods) to odor A and odor B. Pearson’s correlation shown. (**M**) Example cell response to receipt of water. Mean calcium activity at reward outcome of an example cell, shading shows standard error. (**N**) Population responses to reward outcome, all columns sorted by response to water reward on no information trials.

**Fig. S6.**
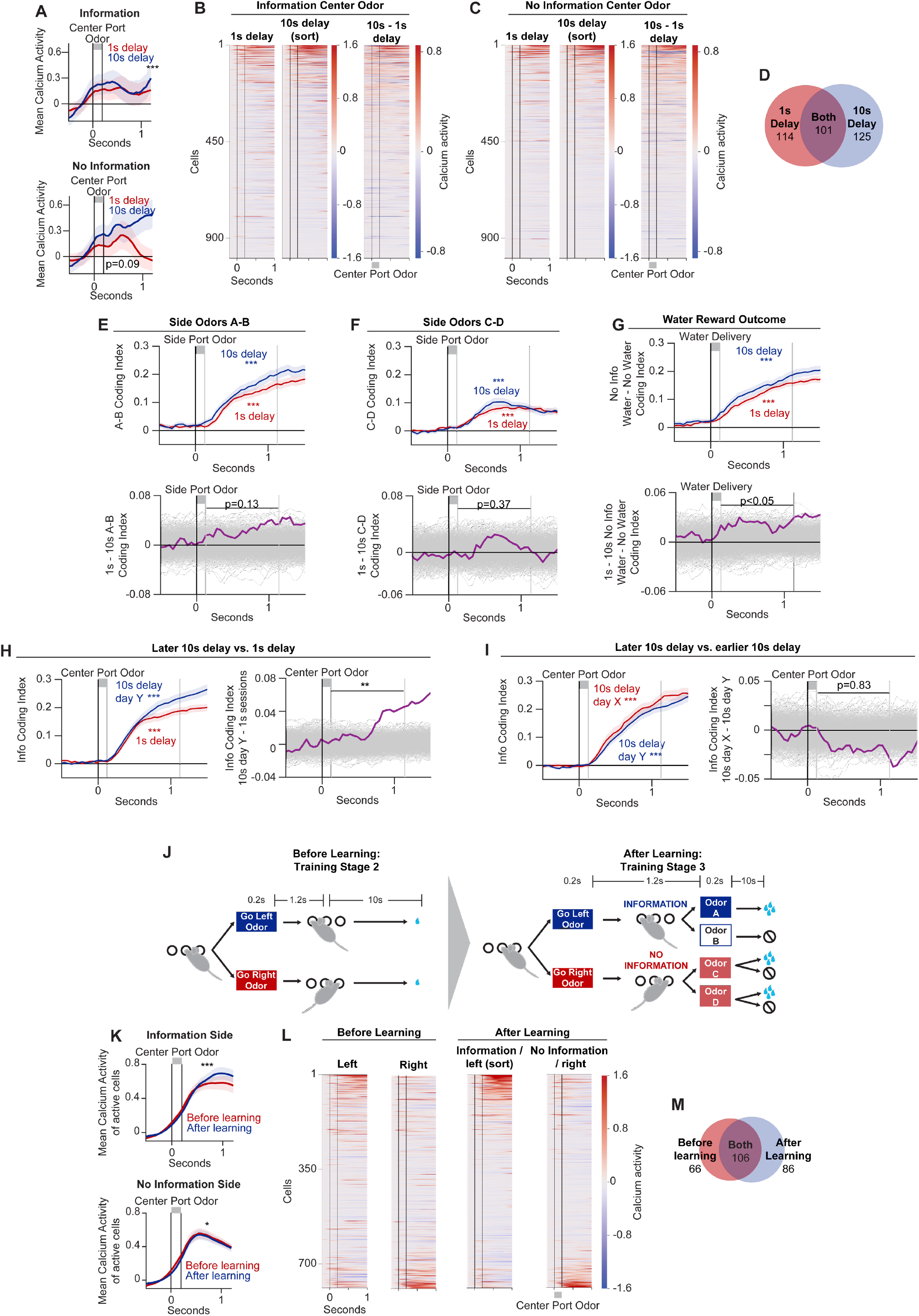
(**A**) Mean population calcium activity at the center port on forced information (top) and forced no information (bottom) trials, in sessions with a 1s (red) and 10s (blue) delay. Shading shows standard error. Significance of the difference between 10s and 1s delay in both top and bottom by signrank test. (**B**) Population activity on forced information trials across delays. Each cell’s mean activity in sessions with 1s and 10s delay, and the difference between them, shown. Cells in all columns sorted by 10s delay activity. (**C**) As in (B), but for forced no information trials. (**D**) Number of cells with significant coding of information at the center port across delays and their overlap, p<0.001, chi-square test, N=993. Coding computed as information index in 1s interval after – 1s interval before the center port odor. Significance (p<0.05) determined by comparison with shuffled data. (**E**) Coding of water CS, side odor A – side odor B, in sessions with 10s vs. 1s delay. Top, mean population coding index for odor A – odor B in 10s and 1s delay sessions. Shading shows standard error, significance of information coding at center odor based on index difference between 1s interval after odor presentation and 1s interval before greater than shuffled data. Bottom, mean difference between coding index in 10s – 1s delay sessions (purple). Gray shows shuffled data. (**F**) Mean population coding index for odor C – odor D in 10s and 1s delay sessions, as in (E). (**G**) Coding of water US, no information water reward – no reward, in sessions with 10s vs. 1s delay as in (E). (**H**) We compared 1s delay sessions with a second separate set of later sessions with the delay returned to 10s. We observed that information coding was diminished in the 1s sessions compared also to these additional later “Day Y” 10s sessions. Left, shows mean population information value coding index in 1s and later 10s delay sessions. Shading shows standard error, significance of information coding at center odor based on index difference between 1s interval after odor presentation and 1s interval before greater than shuffled data. Right, mean difference between information value coding index in later 10s (Day Y) – 1s delay sessions (purple). Gray shows shuffled data. Significance of the difference in coding between 1s and 10s sessions based on observed difference greater than shuffled data. (**I**) We compared the two sets of 10s delay sessions, one earlier before the 1s sessions, and one later after decreasing the delay to 1s and then returning it to 10s. Left, shows mean population information value coding index in earlier Day X 10s and later Day Y 10s delay sessions. Shading shows standard error, significance of information coding at center odor based on index difference between 1s interval after odor presentation and 1s interval before greater than shuffled data. Right, mean difference between information value coding index in earlier 10s (Day X) - later 10s (Day Y) delay sessions (purple). Gray shows shuffled data. Significance of the difference in coding between earlier and later sessions based on observed difference greater than shuffled data. (**J**) Schematic of learning across training stages when information is introduced by providing odor at the side port and making water rewards probabilistic. (**K**) Mean calcium activity of cells responding to the center port odor on forced information (top) and forced no information (bottom) trials, in sessions before (red) and after (blue) learning upon the introduction of information. Shading shows standard error. Significance of difference between before and after learning in both by signrank test. (**L**) Population activity at the center port across learning. Left, activity on forced left (port 1) and forced right (port 2) trials. Right, activity on forced information (port 1) and forced no information (port 2) trials. Each cell’s mean activity shown. Cells in all columns sorted by activity on forced information trials after learning. Information is assigned to either the left or right port across animals. Here, the label “left” indicates whichever port was assigned to provide information, which was counterbalanced across mice. (**M**) Number of cells with significant coding of left vs. right (before learning) and information vs. no information (after learning) at the center port and their overlap, p<0.001, chi-square test, N=788. Coding computed as information index in 1s interval after – 1s interval before the center port odor. Significance (p<0.05) determined by comparison with shuffled data.

**Fig. S7.**
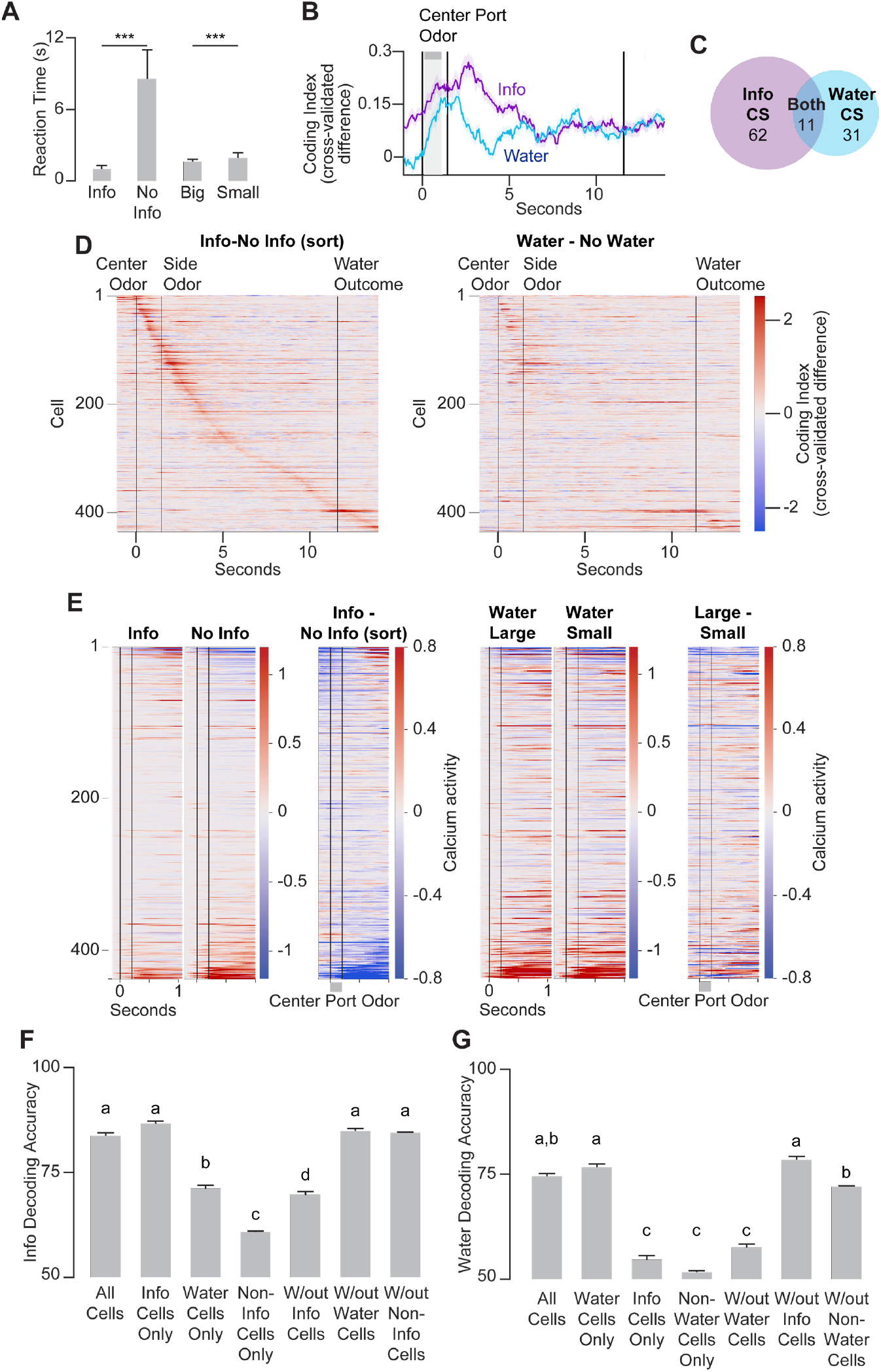
(**A**) Reaction times by trial type in the task variant with four center port odors. Bar shows mean across animals, points show mean value for individual mice. (**B**) Population mean information and water coding indices. Shading shows standard error. Gray vertical shading shows 1s post-center port odor information and water CS coding analysis window. Information coding index compares information and no information trials; water coding index compares large and small water reward trials. (**C**) Number of cells with differential activity (significant coding index increase after odor) for information and water at the center port odor. N=437, overlap, p>0.05, chi-square test. (**D**) Mean information (left) and water (right) coding indices across the population. Cells in both plots sorted by information coding index. (**E**) Population responses to center port odors. Mean calcium activity at the center port on the indicated trial type for each cell and the differences between information and no information and large and small water trials. All cells sorted by the information-no information difference. (**F**) Decoding accuracy of information vs. no information trials 0.8-1s after center odor. Info cells are those with weights in the information-no information classifier trained using all cells in the top 10% of absolute value. Water cells are those in the top 10% of water classifier weight absolute value. Accuracy shown for classifiers trained using all cells, only the top 10% of information classifier cells, only the top 10% of water classifier cells, a matched number of randomly selected other cells, removing only the top 10% of information classifier cells, removing the top 10% of water classifier cells, or removing a matched number of randomly selected other cells. Significant differences (labels a-d) tested with kruskal-wallis test with tukey-kramer correction for multiple comparisons. (**G**) Decoding accuracy of large vs. small water reward trials 0.8-1s after center odor. Accuracy shown for classifiers trained using all cells, only the top 10% of water classifier cells, only the top 10% of info classifier cells, a matched number of randomly selected other cells, removing only the top 10% of water classifier cells, removing the top 10% of info classifier cells, or removing a matched number of randomly selected other cells. Significant differences (labels a-c) tested with kruskal-wallis test with tukey-kramer correction for multiple comparisons.

**Fig. S8.**
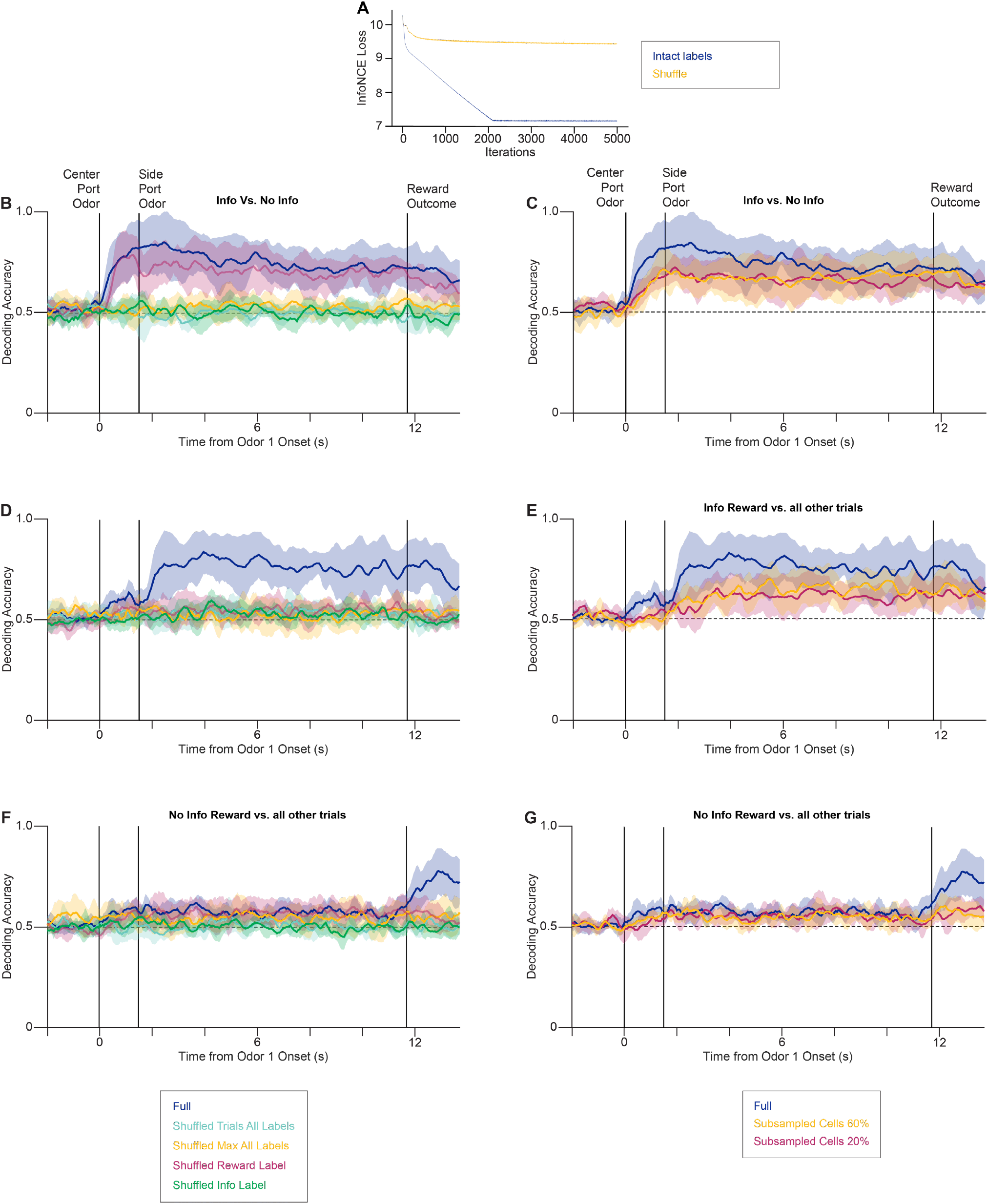
(**A**) Comparison of the model’s contrastive loss as measured by InfoNCE (Info Noise-Contrastive Estimation)(*80*) with intact versus shuffled labels. Lower loss suggests a better fit model. Decoding accuracy of (**B**) information trial type versus no information trial type for all trials, (**C**) presence of reward on information trials and (**D**) presence of reward on no information trials. In (b-d), the legends are defined as follows. In the “shuffled trials all labels” condition, the assignment of trials was randomized relative to their original time-varying labels, for all the label types, but we left the matching between labels and the time they were assigned in a trial (table S3) intact. In the “shuffled max all labels” condition, we randomly scrambled both the trial- and the time-affiliation of a given label, for all label types, thus resulting in the maximum amount of signal scrambling. For the “shuffled info label” condition, we only scrambled the trial-label assignment for the “info vs no info” label type (table S3), leaving all others intact. For the “shuffled reward label” condition, we only scrambled the trial-label assignment for the “rewarded vs unrewarded” label (table S3), leaving all others intact. “Full” is the finalized intact labels for the full model. Results are reported as an average across 5 runs, using different random seeds to achieve the train/test allocation. (**E-G**) Models fit to only random sub-populations of cells were also used as controls to show that embedding quality deteriorated with reduced sample size. 20% (magenta) and 60% (orange) of OFC neurons recorded were sub-sampled and compared to the full sample (blue). Each sampling was the average of five runs, using five different random seeds to subsample the cells. Results are reported averaged across sessions, animals and runs with error bars to show 95% confidence interval.

**Table S1.**
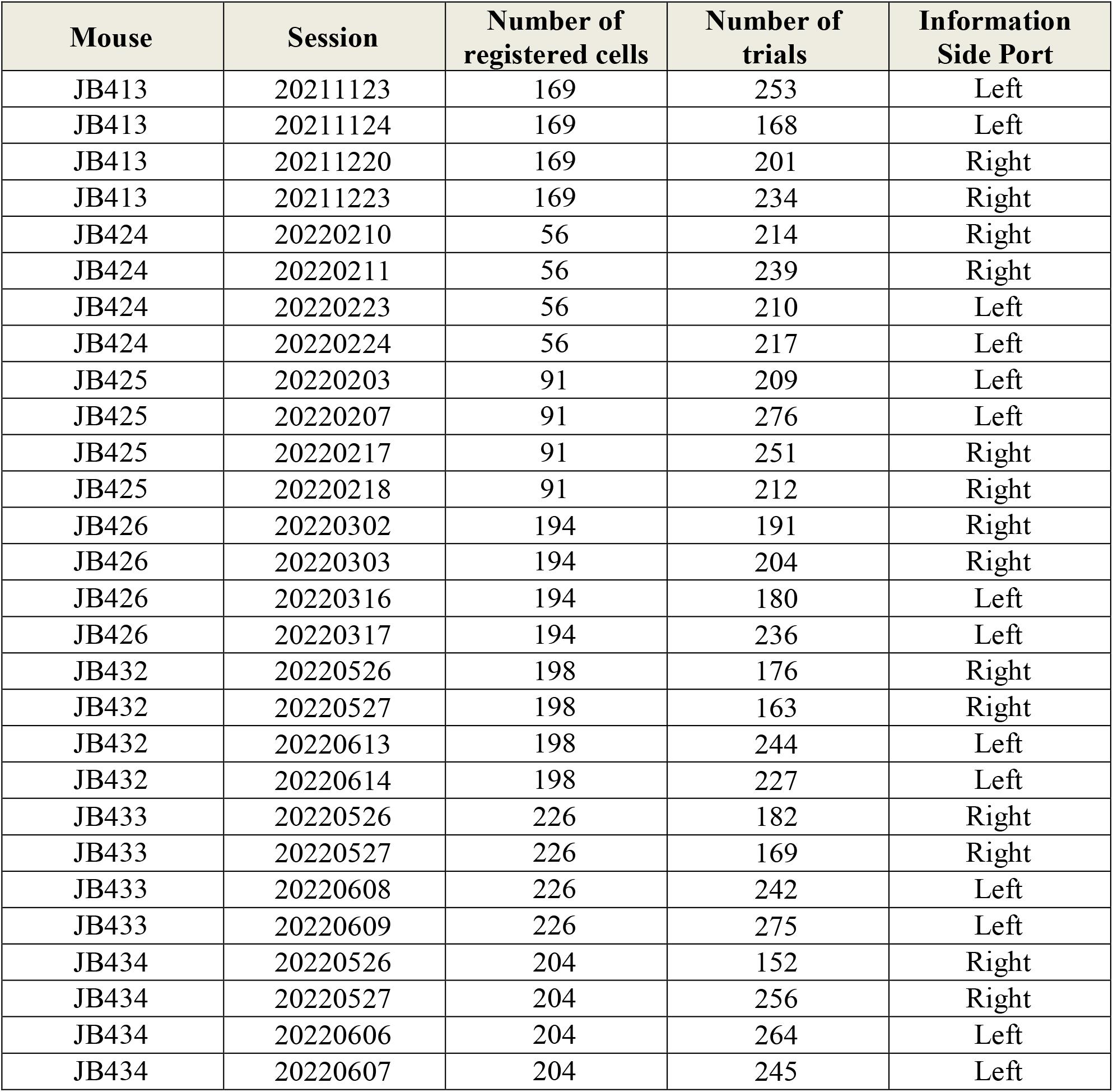
Mice and sessions imaged in main dataset.

**Table S2.** Mice and sessions imaged in the dataset comprising learning before and after the introduction of information.

| Mouse | Session | Number of registered cells | Number of trials | Pre vs. Post Learning |
| --- | --- | --- | --- | --- |
| JB413 | '20211014' | 148 | 102 | Pre |
| JB413 | '20211015' | 148 | 159 | Pre |
| JB413 | '20211102' | 148 | 196 | Post |
| JB413 | '20211103' | 148 | 197 | Post |
| JB424 | '20220105' | 197 | 210 | Pre |
| JB424 | '20220106' | 197 | 254 | Pre |
| JB424 | '20220121' | 197 | 218 | Post |
| JB424 | '20220124' | 197 | 227 | Post |
| JB425 | '20220105' | 48 | 222 | Pre |
| JB425 | '20220106' | 48 | 310 | Pre |
| JB425 | '20220119' | 48 | 218 | Post |
| JB425 | '20220120' | 48 | 227 | Post |
| JB426 | '20220126' | 152 | 218 | Pre |
| JB426 | '20220127' | 152 | 210 | Pre |
| JB426 | '20220131' | 152 | 252 | Post |
| JB426 | '20220201' | 152 | 231 | Post |
| JB432 | '20220428' | 77 | 185 | Pre |
| JB432 | '20220429' | 77 | 179 | Pre |
| JB432 | '20220505' | 77 | 203 | Post |
| JB432 | '20220506' | 77 | 166 | Post |
| JB433 | '20220428' | 106 | 164 | Pre |
| JB433 | '20220429' | 106 | 156 | Pre |
| JB433 | '20220505' | 106 | 194 | Post |
| JB433 | '20220506' | 106 | 191 | Post |
| JB434 | '20220428' | 60 | 218 | Pre |
| JB434 | '20220429' | 60 | 210 | Pre |
| JB434 | '20220504' | 60 | 252 | Post |
| JB434 | '20220505' | 60 | 231 | Post |

**Table S3.** Mice and sessions imaged in delay comparison sessions, 10s vs. 1s.

| Mouse | Session | Number of registered cells | Number of trials | 10s vs 1s |
| --- | --- | --- | --- | --- |
| JB424 | '20220527' | 169 | 128 | 10 |
| JB424 | '20220531' | 169 | 165 | 10 |
| JB424 | '20220606' | 169 | 212 | 1 |
| JB424 | '20220607' | 169 | 142 | 1 |
| JB425 | '20220518' | 135 | 170 | 10 |
| JB425 | '20220519' | 135 | 135 | 10 |
| JB425 | '20220607' | 135 | 133 | 1 |
| JB425 | '20220609' | 135 | 174 | 1 |
| JB426 | '20220616' | 72 | 213 | 10 |
| JB426 | '20220617' | 72 | 244 | 10 |
| JB426 | '20220711' | 72 | 297 | 1 |
| JB426 | '20220713' | 72 | 261 | 1 |
| JB432 | '20220623' | 258 | 200 | 10 |
| JB432 | '20220627' | 258 | 169 | 10 |
| JB432 | '20220706' | 258 | 279 | 1 |
| JB432 | '20220707' | 258 | 217 | 1 |
| JB433 | '20220629' | 136 | 246 | 1 |
| JB433 | '20220630' | 136 | 230 | 1 |
| JB433 | '20220708' | 136 | 207 | 10 |
| JB433 | '20220711' | 136 | 207 | 10 |
| JB434 | '20220812' | 223 | 220 | 10 |
| JB434 | '20220817' | 223 | 178 | 10 |
| JB434 | '20220824' | 223 | 447 | 1 |
| JB434 | '20220825' | 223 | 418 | 1 |

**Table S4.**
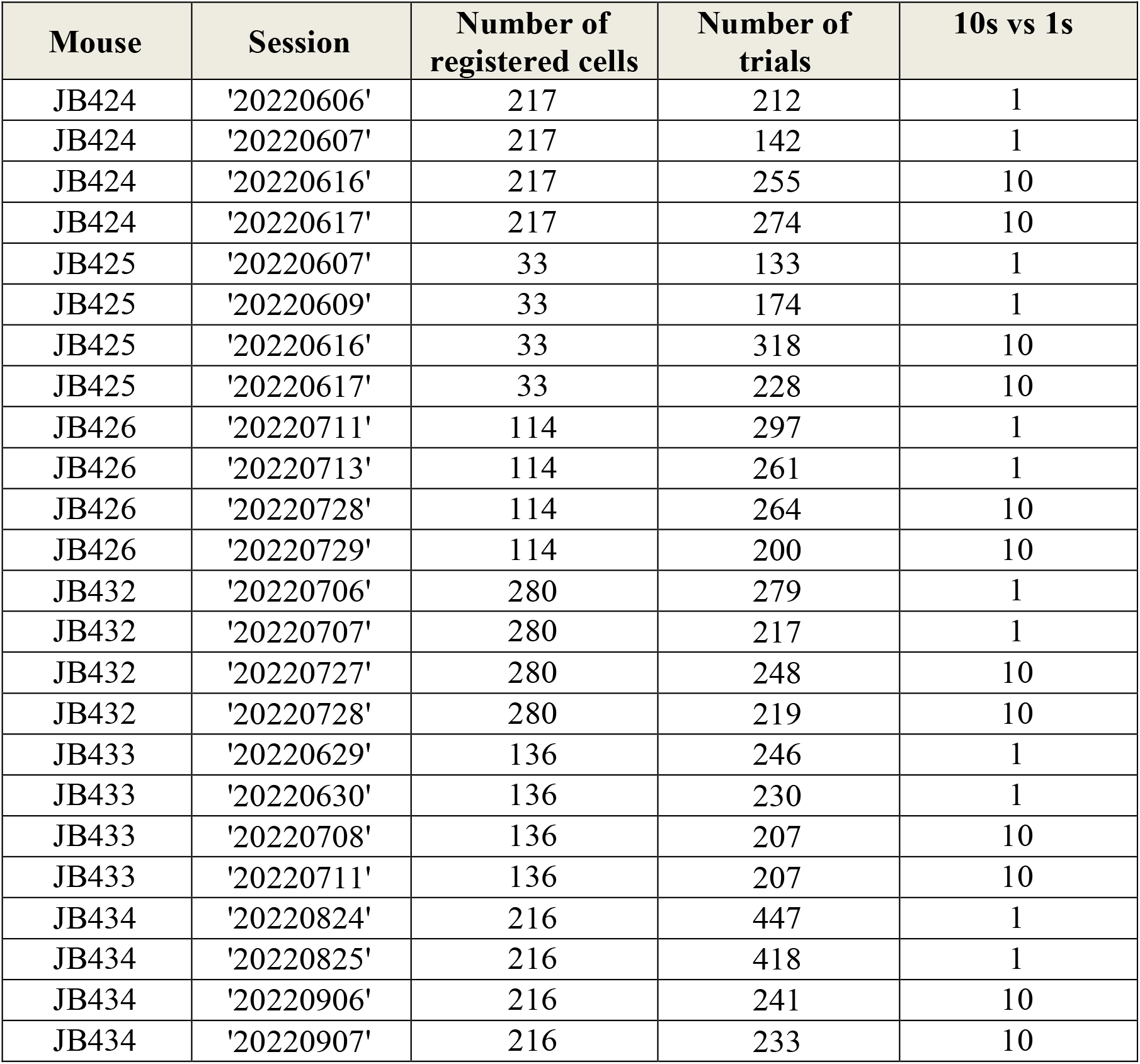
Mice and sessions imaged in analyses comparing 1s sessions to later 10s sessions.

**Table S5.** Mice and sessions imaged in early vs. late 10s sessions.

| **Mouse** | **Session** | **Number of**<br>**registered cells** | **Number of**<br>**trials** | **10s vs 1s** |
| --- | --- | --- | --- | --- |
| JB424 | '20220527' | 152 | 128 | 10 |
| JB424 | '20220531' | 152 | 165 | 10 |
| JB424 | '20220616' | 152 | 255 | 10 |
| JB424 | '20220617' | 152 | 274 | 10 |
| JB425 | '20220518' | 34 | 170 | 10 |
| JB425 | '20220519' | 34 | 135 | 10 |
| JB425 | '20220616' | 34 | 318 | 10 |
| JB425 | '20220617' | 34 | 228 | 10 |
| JB426 | '20220616' | 121 | 213 | 10 |
| JB426 | '20220617' | 121 | 244 | 10 |
| JB426 | '20220728' | 121 | 264 | 10 |
| JB426 | '20220729' | 121 | 200 | 10 |
| JB432 | '20220623' | 139 | 200 | 10 |
| JB432 | '20220627' | 139 | 169 | 10 |
| JB432 | '20220727' | 139 | 248 | 10 |
| JB432 | '20220728' | 139 | 219 | 10 |
| JB433 | '20220621' | 161 | 185 | 10 |
| JB433 | '20220711' | 161 | 207 | 10 |
| JB434 | '20220812' | 220 | 220 | 10 |
| JB434 | '20220817' | 220 | 178 | 10 |
| JB434 | '20220906' | 220 | 241 | 10 |
| JB434 | '20220907' | 220 | 233 | 10 |

**Table S6.** Cells per session for *full* CEBRA model.

| Mouse | Session | Number of cells | Number of trials (per session) |
| --- | --- | --- | --- |
| JB432 | 20220526 | 442 | 56 |
| JB432 | 20220527 | 522 | 44 |
| JB432 | 20220613 | 505 | 68 |
| JB432 | 20220614 | 426 | 54 |
| JB426 | 20220302 | 314 | 48 |
| JB426 | 20220303 | 599 | 52 |
| JB426 | 20220316 | 381 | 32 |
| JB426 | 20220317 | 312 | 56 |
| JB434 | 20220526 | 417 | 24 |
| JB434 | 20220527 | 323 | 60 |
| JB434 | 20220606 | 547 | 16 |
| JB434 | 20220607 | 516 | 64 |
| JB413 | 20211123 | 331 | 48 |
| JB413 | 20211124 | 475 | 24 |
| JB413 | 20211220 | 397 | 44 |
| JB413 | 20211223 | 334 | 68 |
| JB424 | 20220210 | 414 | 48 |
| JB424 | 20220211 | 337 | 32 |
| JB424 | 20220223 | 416 | 24 |
| JB424 | 20220224 | 358 | 32 |
| JB425 | 20220203 | 292 | 36 |
| JB425 | 20220207 | 341 | 36 |
| JB425 | 20220217 | 177 | 48 |
| JB425 | 20220218 | 179 | 56 |

**Table S7.** Cells per session for *delay* CEBRA model.

| Mouse | Session | Number of cells | Number of trials |
| --- | --- | --- | --- |
| JB432 | 20220623 | 258 | 44 |
| JB432 | 20220627 | 258 | 40 |
| JB432 | 20220706 | 258 | 92 |
| JB432 | 20220707 | 258 | 56 |
| JB433 | 20220629 | 136 | 64 |
| JB433 | 20220630 | 136 | 72 |
| JB433 | 20220708 | 136 | 64 |
| JB433 | 20220711 | 136 | 64 |
| JB426 | 20220616 | 72 | 56 |
| JB426 | 20220617 | 72 | 44 |
| JB426 | 20220711 | 72 | 92 |
| JB426 | 20220713 | 72 | 88 |
| JB434 | 20220812 | 223 | 36 |
| JB434 | 20220817 | 223 | 24 |
| JB434 | 20220824 | 223 | 203* |
| JB434 | 20220825 | 223 | 84 |
| JB424 | 20220527 | 169 | 48 |
| JB424 | 20220531 | 169 | 80 |
| JB424 | 20220606 | 169 | 120 |
| JB424 | 20220607 | 169 | 80 |
| JB425 | 20220518 | 135 | 84 |
| JB425 | 20220519 | 135 | 68 |
| JB425 | 20220607 | 135 | 80 |
| JB425 | 20220609 | 135 | 88 |

**Table S8.** CEBRA labels.

| **Label Type** | **Range** | **Type** | **Start Frame** | **End Frame** |
| --- | --- | --- | --- | --- |
| trial time index | [1, 2,3,...,n_time] | discrete | 1 | n_time |
| rewarded vs unrewarded | [-1, 0, 1] | discrete | 274 | n_time |
| info vs no info | [-1, 0, 1] | discrete | 40 | n_time |
| info rewarded vs unrewarded | [0, 1, 2] | discrete | 70 | n_time |
| no info rewarded vs unrewarded | [-2, -1, 0] | discrete | 274 | n_time |
| trial epoch center port odor | [0, 1] | discrete | 40 | 44 |
| trial epoch side port odor | [0, 2] | discrete | 70 | 74 |
| trial epoch outcome | [0, 3] | discrete | 74 | n_time |

**Table S9.** Full behavioral model parameters for each animal cross-animal parameter mean and SEM. See Methods for behavior model details.

| Mouse | $\sigma$ | $\beta$ | $\alpha 1$ | $\alpha 2$ | InfoVal | $\lambda$ | E |
| --- | --- | --- | --- | --- | --- | --- | --- |
| 1 | 0.244 | 0.992 | 0.302 | 0.200 | 1 | N/A | -0.258 |
| 2 | 0.335 | 0.634 | 0.300 | 0.208 | 1 | N/A | -0.236 |
| 3 | 0.235 | 0.600 | 0.312 | 0.250 | 1 | N/A | -0.261 |
| 4 | 0.370 | 0.865 | 0.302 | 0.237 | 1 | N/A | -0.267 |
| 5 | 0.303 | 0.600 | 0.733 | 0.433 | 1 | 2.000 | -0.232 |
| 6 | 0.350 | 0.600 | 0.960 | 0.459 | 1 | 1.875 | -0.291 |
| 7 | 0.235 | 0.934 | 0.303 | 0.463 | 1 | 0.409 | -0.252 |
| 8 | 0.227 | 0.611 | 0.461 | 0.106 | 1 | 0.300 | -0.271 |
| 9 | 0.269 | 0.824 | 0.484 | 0.153 | 1 | 0.653 | -0.225 |
| 10 | 0.399 | 0.619 | 0.829 | 0.061 | 1 | 0.323 | -0.232 |
| **Mean** | 0.297 | 0.728 | 0.499 | 0.257 | 1 | 0.927 | -0.253 |
| **SEM** | 0.020 | 0.050 | 0.251 | 0.146 | N/A | 0.794 | 0.021 |

**Table S10.**
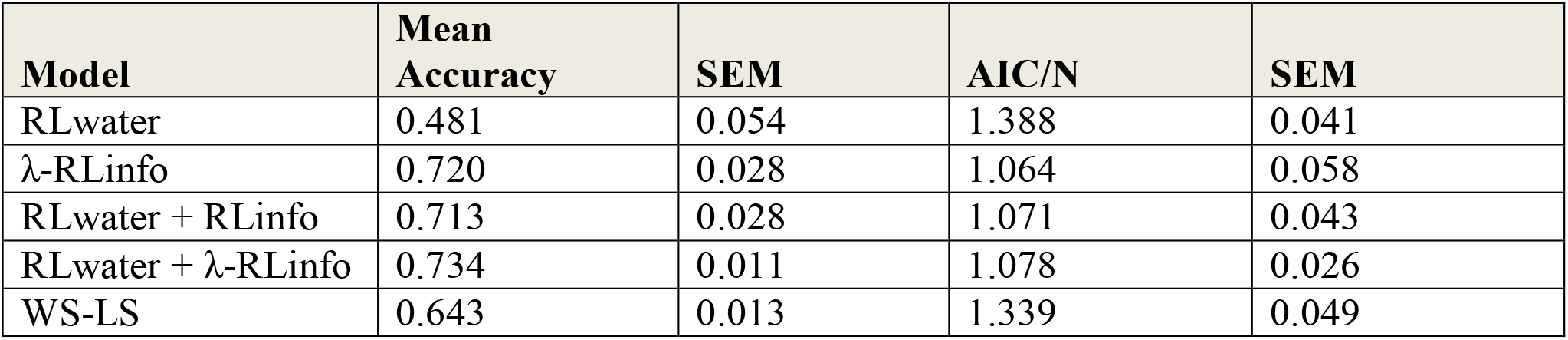
Behavioral model comparison. Accuracy (SEM) and Aikake Information Criterion normalized by sample size, N, (SEM) reported across animals for each tested behavioral model. See Methods for behavioral model details.

**Table S11.** Summary statistics for test-train splits in behavioral model fitting. Accuracy (SEM) and Aikake Information Criterion normalized by sample size, N, (SEM) reported across animals for test-train splits in behavioral models.

| **Model** | **Test Accuracy** | **SEM** | **Train AIC/N** | **SEM** |
| --- | --- | --- | --- | --- |
| RLwater | 0.483 | 0.014 | 1.39 | 0.01 |
| λ-RLinfo | 0.719 | 0.009 | 1.07 | 0.01 |
| RLwater + RLinfo | 0.715 | 0.012 | 1.08 | 0.01 |
| RLwater + λ-RLinfo | 0.74 | 0.014 | 1.07 | 0.02 |
| WS-LS | 0.646 | 0.036 | 1.31 | 0.03 |

